# Decoding Estimates of Curvilinear Self-Motion from Neural Signals in a Model of Primate MSTd

**DOI:** 10.1101/2021.09.29.462288

**Authors:** Oliver W. Layton, Nathaniel Powell, Scott T Steinmetz, Brett R Fajen

## Abstract

Self-motion produces characteristic patterns of optic flow on the eye of the mobile observer. Movement along linear, straight paths without eye movements yields motion that radiates from the direction of travel (heading). The observer experiences more complex motion patterns while moving along more general curvilinear (e.g. circular) paths, the appearance of which depends on the radius of the curved path (path curvature) and the direction of gaze. Neurons in brain area MSTd of primate visual cortex exhibit tuning to radial motion patterns and have been linked with linear heading perception. MSTd also contains neurons that exhibit tuning to spirals, but their function is not well understood. We investigated in a computational model whether MSTd, through its diverse pattern tuning, could support estimation of a broader range of self-motion parameters from optic flow than has been previously demonstrated. We used deep learning to decode these parameters from signals produced by neurons tuned to radial expansion, spiral, ground flow, and other patterns in a mechanistic neural model of MSTd. Specifically, we found that we could accurately decode the clockwise/counterclockwise sign of curvilinear path and the gaze direction relative to the path tangent from spiral cells; heading from radial cells; and the curvature (radius) of the curvilinear path from activation produced by both radial and spiral populations. We demonstrate accurate decoding of these linear and curvilinear self-motion parameters in both synthetic and naturalistic videos of simulated self-motion. Estimates remained stable over time, while also rapidly adapting to dynamic changes in the observer’s curvilinear self-motion. Our findings suggest that specific populations of neurons in MSTd could effectively signal important aspects of the observer’s linear and curvilinear self-motion.

**Author Summary:** How do we perceive our self-motion as we move through the world? Substantial evidence indicates that brain area MSTd contains neurons that signal the direction of travel during movement along straight paths. We wondered whether MSTd neurons could also estimate more general self-motion along curved paths. We tested this idea by using deep learning to decode signals produced by a neural model of MSTd. The system accurately decoded parameters that specify the observer’s self-motion along straight and curved paths in videos of synthetic and naturalistic scenes rendered in the Unreal game engine. Our findings suggest that MSTd could jointly signal self-motion along straight and curved paths.

## Introduction

Substantial progress has been made over the past several decades toward understanding how neurons in visual cortex encode information about the environment and state of the observer. The pioneering work of Hubel & Wiesel (Hubel, 1982) exemplifies the successes that have been garnered in primary visual cortex and other areas early in the visual hierarchy (Hubel & Wiesel, 2012). Tuning properties and the function of neurons in areas further along in extrastriate cortex, however, have remained more elusive.

One such extrastriate area, the dorsal medial superior temporal area (MSTd), has long been implicated in encoding information that subserves self-motion perception. Early studies demonstrated tuning to diverse motion patterns that encompass large portions of the visual field, such as spirals (Graziano, Andersen, & Snowden, 1994) and radial expansion (Saito et al., 1986) (Duffy & Wurtz, 1991) (Duffy & Wurtz, 1995). Sensitivity to radial expansion is notable because it resembles the pattern of optic flow experienced during linear translation: movement along a straight path (Figure 1a) without eye movements (Gibson, 1950). The motion radiates from a singularity known as the focus of expansion (FoE) that corresponds to the direction of self-motion (heading). Translation, together with another component known as rotation, fully characterize the possible first-order optic flow patterns that may arise by an observer moving through a rigid environment (Longuet-Higgins & Prazdny, 1980). Rotation arises during eye movements or self-motion along a curvilinear path (Figure 1b) (Warren, Mestre, Blackwell, & Morris, 1991) (Raudies & Neumann, 2013). Findings from the past several decades generally support the hypothesis that MSTd encodes information relating to translational (Duffy & Wurtz, 1995) (Logan & Duffy, 2006) (Gu, Watkins, Angelaki, & DeAngelis, 2006) (Gu, DeAngelis, & Angelaki, 2012) (Britten & van Wezel, 1998) (Britten & van Wezel, 2002) and rotational (Shenoy, Bradley, & Andersen, 1999; Page & Duffy, 1999; Komatsu & Wurtz, 1988) optic flow, though the nature of the representation remains unclear (Orban et al., 1992; Danz, Angelaki, & DeAngelis, 2020).

**Figure 1:**
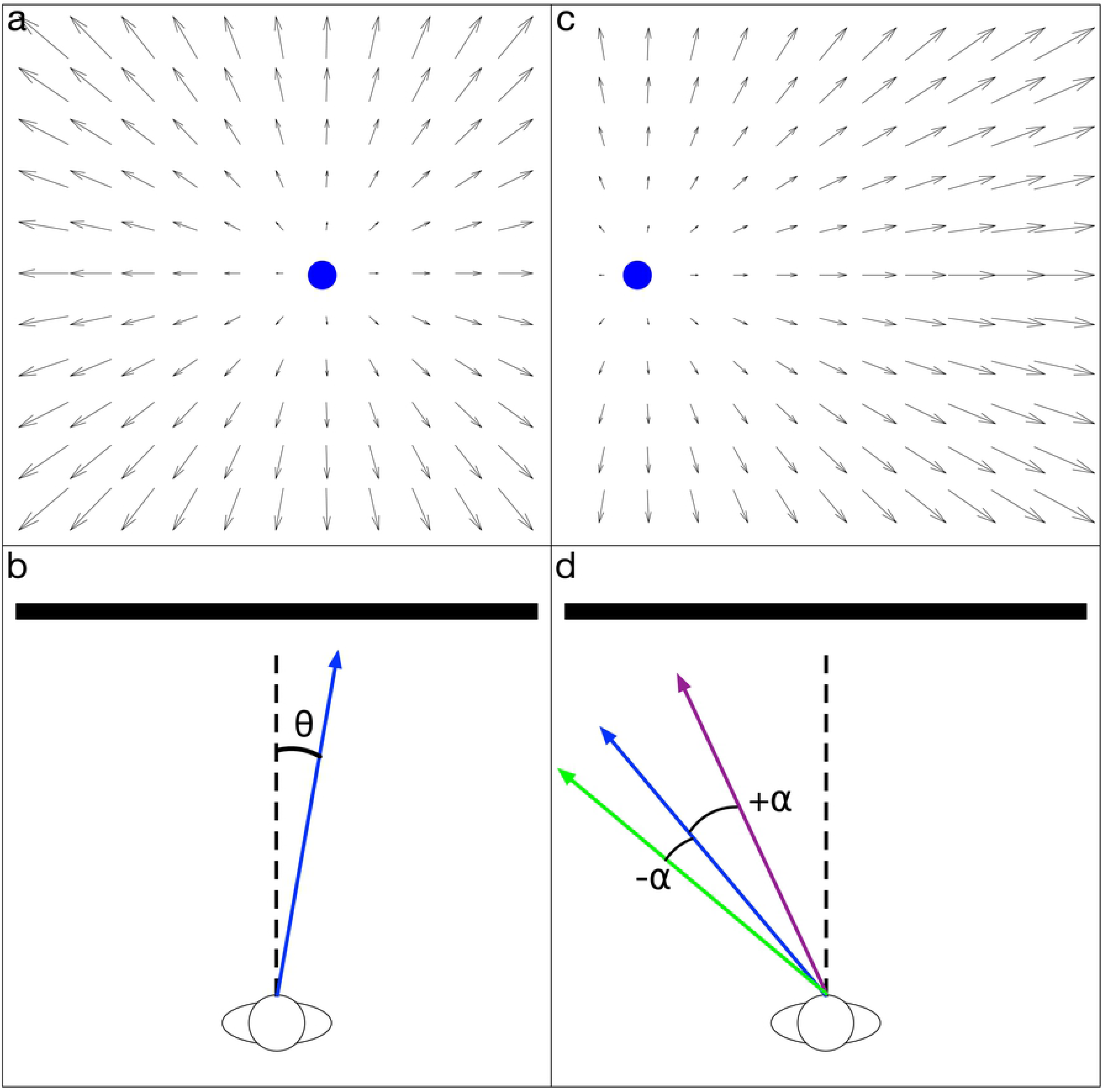
Optic flow fields generated by different types of linear and curvilinear self-motion over a ground plane. (A) Self-motion along straight-ahead (left) and right-ward (right) linear trajectories. (B) Curvilinear self-motion along circular paths with different radii. (C) Curvilinear self-motion along CW (left) and CCW (right) 5 m radius circular paths. (D) Curvilinear self-motion along 5 m radius circular paths with the observer gaze offset from the path tangent (left: gaze directed outside the future path, center: gaze along path tangent as in B-C; right: gaze directed inside the future path).

One perspective that may offer insight about the underlying representation that has received less attention is decoding the information carried by the neural signals. The central idea is that an algorithm maps the neural activity elicited by different stimuli (e.g. optic flow) to descriptive parameters (e.g. heading direction). Accurately decoding such parameters would suggest that a neural population may signal these stimulus parameters, perhaps limiting the precision with which they are perceived (Gu, Fetsch, Adeyemo, Deangelis, & Angelaki, 2010). Computational models have had success in explaining human judgments by decoding heading from model MSTd signals using the direction preference of the maximally active neuron (Royden, 2002) (Layton, Mingolla, & Browning, 2012) (Browning, Grossberg, & Mingolla, 2009). However, this simplistic winner-take-all approach to decoding is limited by the assumption that individual neurons exhibit one-to-one correspondence to the decoded stimulus parameter (e.g. heading). It fails to account for data suggesting that heading estimates rely on representations distributed across multiple MSTd subpopulations (Xu, Wallisch, & Bradley, 2014). Challenges also arise when decoding multiple stimulus parameters (e.g. heading and curvilinear path) since a single neuron would need to demonstrate tuning to the ideal combination of parameter values, leading to combinatorial explosion. Population vector decoding (Georgopoulos, Schwartz, & Kettner, 1986) addresses these shortcomings, but the approach performs poorly when applied to neural data from MSTd compared to more complex techniques (Gu et al., 2010), such as maximum likelihood estimation (Jazayeri & Movshon, 2006) (Sanger, 1996).

Here we present an alternative approach that uses deep learning to decode self-motion from a mechanistic model of MSTd that processes optic flow. We simulated the competitive dynamics (CD) model, a neural model that contains units that emulate physiological properties of the medial temporal area (MT) and MSTd and accounts for a range of human behavioral data on self-motion perception (Layton & Browning, 2012) (Layton et al., 2012) (Layton & Fajen, 2016b) (Layton & Fajen, 2017). We used the neural model and decoder system to explore the possibility that MSTd could represent not only heading during linear self-motion, as has been well-established (Gu et al., 2012) (Duffy & Wurtz, 1995) (Britten & van Wezel, 2002), but also curvilinear self-motion (Figure 1b-d) whose neural basis is presently unknown. We demonstrate accurate decoding of:

- heading when the observer moves along a linear path (henceforth “heading”) from model neurons tuned to radial expansion (henceforth “radial cells”) (Figure 1a).
- curvature when the observer moves along a curvilinear path (henceforth “path curvature”) from radial cells and cells tuned to spiral patterns (henceforth “spiral cells”) (Figure 1b).
- whether the observer moves along a clockwise (CW) or counterclockwise (CCW) curvilinear path (henceforth “path sign”) from spiral cells (Figure 1c).
- the horizontal offset in the observer’s gaze relative to the straight-ahead when the observer moves along a curvilinear path (henceforth “gaze offset”) from spiral cells (Figure 1d).

Simulations of the CD model informed the selection of the above relationships between cell populations and self-motion parameters. The exception is the association between MSTd units tuned to radial expansion and heading perception during linear self-motion, which has long been supported by computational studies (Perrone, 1992) (Royden, 2002) (Layton & Fajen, 2016b). For horizontal gaze offset angle, simulations revealed systematic shifts in the activation of spiral cells (Figure S1b). This association is plausible given that the gaze offset angle modulates the apparent curvature of the flow field: gaze directed outside the future path makes the flow appear more linear and gaze directed inside the future path makes the flow appear more curved (Layton & Browning, 2014) (Li & Cheng, 2011). Moreover, we suspected that cells tuned to opposite spiral orientations may be capable of discriminating the sign of the curvilinear path from the corresponding CW/CCW optic flow patterns (Figure 1c). For path curvature, simulations of the CD model revealed proportional bias in radial cells that signal the observer’s heading direction during linear self-motion (Figure S1c). For example, radial cells tuned to peripheral headings yield greater activation to higher curvature paths than to low curvature paths. Radial cells alone, however, cannot accurately encode path curvature because of the profound influence of that gaze offset angle exerts on optic flow pattern (Figure 1d). If MSTd could accurately account for the gaze offset angle (e.g. via spiral cells), however, perhaps path curvature could be accurately estimated. This led us to consider the possibility that path curvature could be jointly encoded by radial and spiral motion MSTd subpopulations.

We tested the robustness with which the deep network decoder estimates each of these parameters during simulated self-motion through both random dot-textured and more visually realistic outdoor scenes. We demonstrate stable decoding of curvilinear self-motion from the model MSTd signals over time, even as the path curvature changes dramatically throughout the trial. The accuracy of decoded estimates compares favorably to an existing mathematical model (Perrone, 2018) and human data (Li & Cheng, 2011). Our computational investigation demonstrates the feasibility of decoding self-motion parameters other than heading along a linear trajectory from MSTd. It also characterizes the precision with which specific subpopulations in MSTd could represent certain parameters describing the observer’s linear and curvilinear self-motion.

## Results

### Decoding paradigm

Our approach uses deep learning to learn the mapping between MSTd signals generated by the CD neural to model self-motion parameters, if such a relationship exists. Figure 2a summarizes the paradigm that we employed to derive self-motion estimates. We simulated the neural model of MT and MSTd with optic flow produced from either linear or curvilinear self-motion over a synthetic ground plane (dot ground dataset; Figure 2b) and through a rich neighborhood scene created using the AirSim drone simulator (neighborhood dataset; Figure 2c). We employed a supervised learning paradigm wherein neural signals derived from the MSTd activation to each video training set represented the features that we used to train the deep network decoders. The accuracy of decoding was determined using videos of novel self-motion trajectories not encountered during training. The test set consisted of random heading directions for linear self-motion samples and random pairings of path curvature and gaze offset for curvilinear self-motion samples, counterbalancing the CW/CCW path sign. Given the prevalence of ground flow in these datasets, we decoded path curvature and heading using units tuned to ground rather than full-field radial expansion. The activation of units tuned to all the patterns that we simulated contributed to decoding path sign and gaze offset angle.

**Figure 2:**
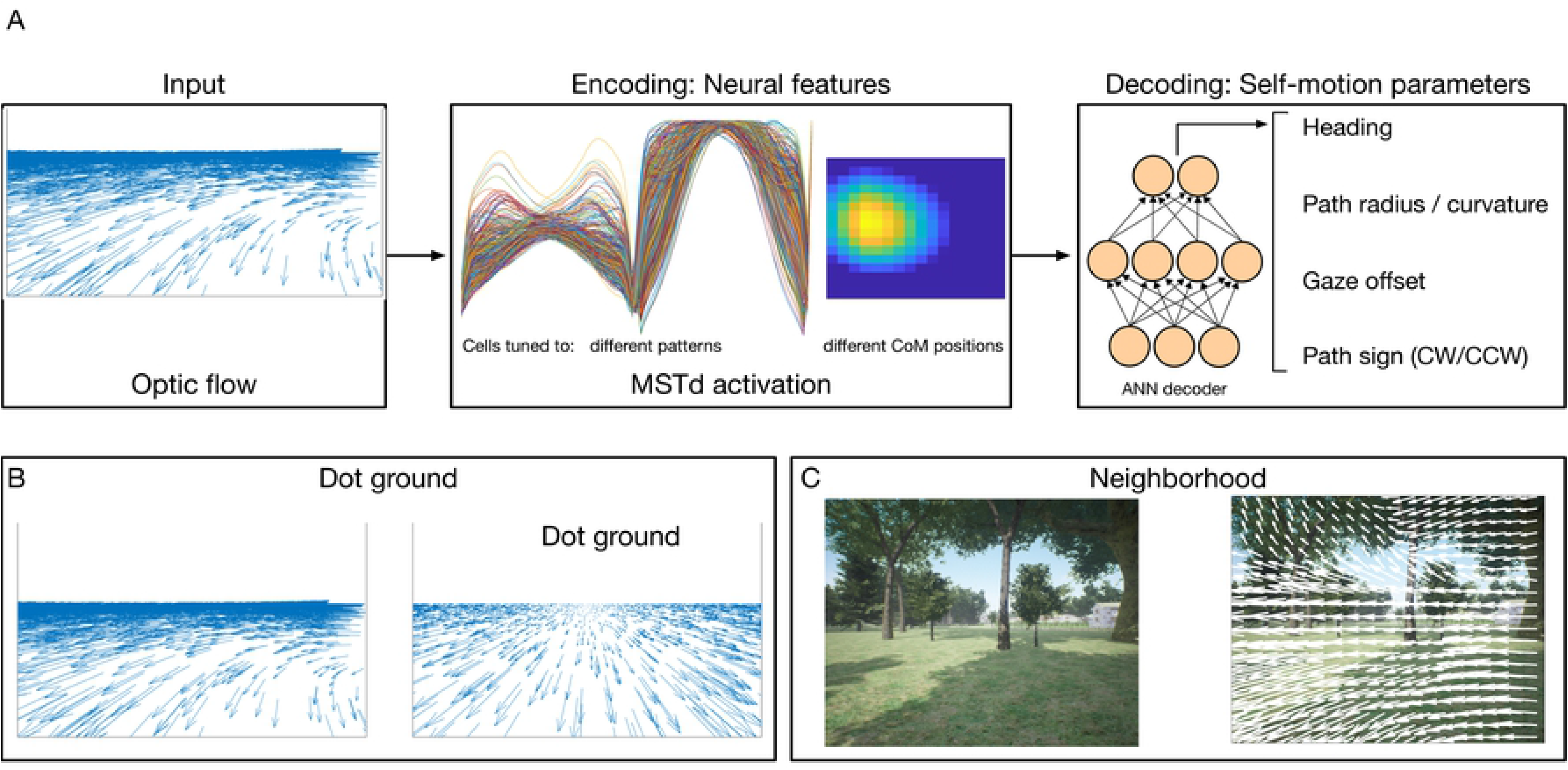
Overview of the neural encoder-decoder framework (A) and sample optic flow fields from the dot ground (B) and neighborhood (C) datasets. (A) Optic flow generated by simulated self-motion is passed through a neural model of areas MT and MSTd, which contains units tuned to radial, ground, spiral, center, and other flow patterns with different preferred center of motion (CoM) locations. The activation of units in MSTd serves as the input to deep networks that decode parameters describing the observer’s self-motion (heading, path curvature, gaze offset, and path sign). (B) Example curvilinear (left) and linear (right) optic flow fields from the dot ground dataset. (C) The neighborhood dataset consists of optic flow detected from videos of self-motion through a neighborhood scene created using the AirSim drone simulator.

### Decoding linear and curvilinear self-motion from model MSTd

We start by investigating the accuracy with which we can decode path sign, gaze offset angle (Figure 3), path curvature (Figure 4-5), and heading (Figure 6) from the hypothesized MSTd cell populations. Here we focus on decoding from a single frame of video (Frame 10) and the subsequent section characterizes the stability of estimates over time.

**Figure 3:**
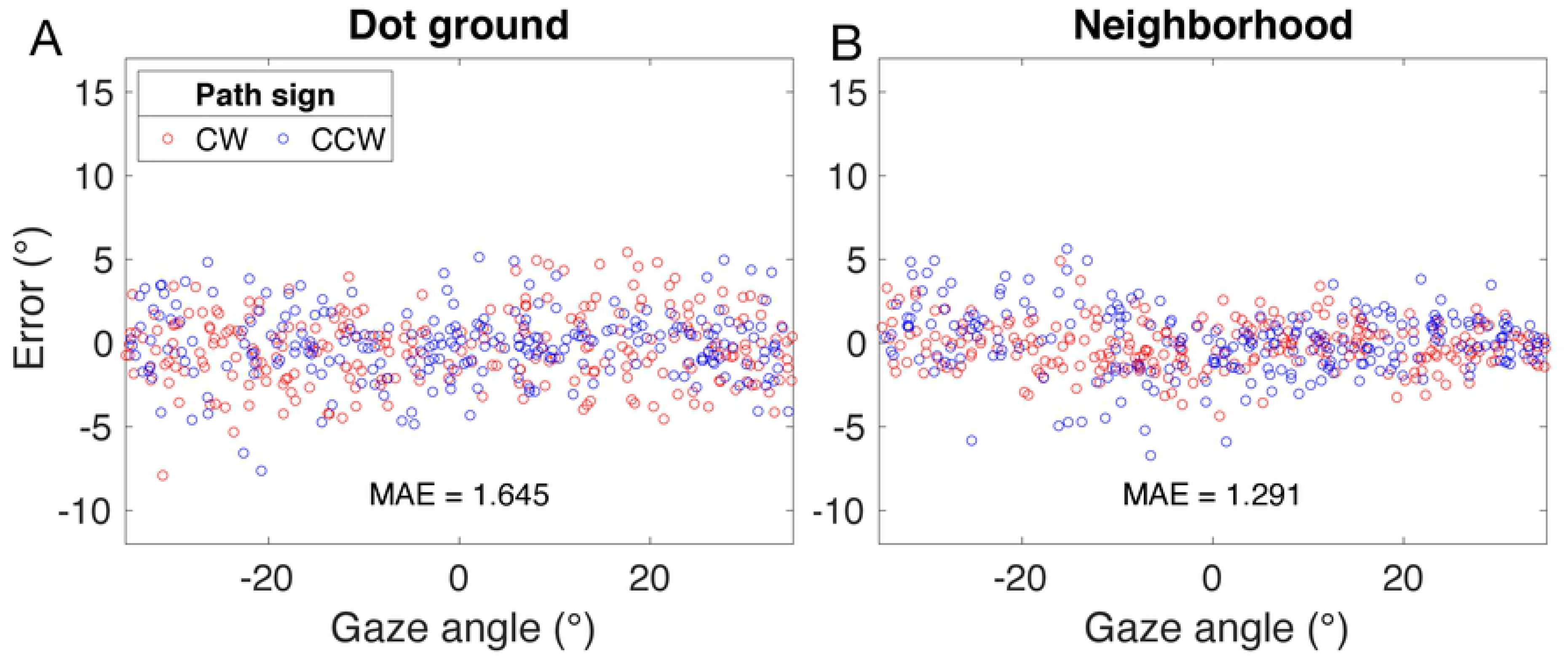
Error in gaze offset estimated from the dot ground (left) and neighborhood (right) test sets. Errors for CW curvilinear paths are shown in red and those for CCW paths are shown in blue.

**Figure 4:**
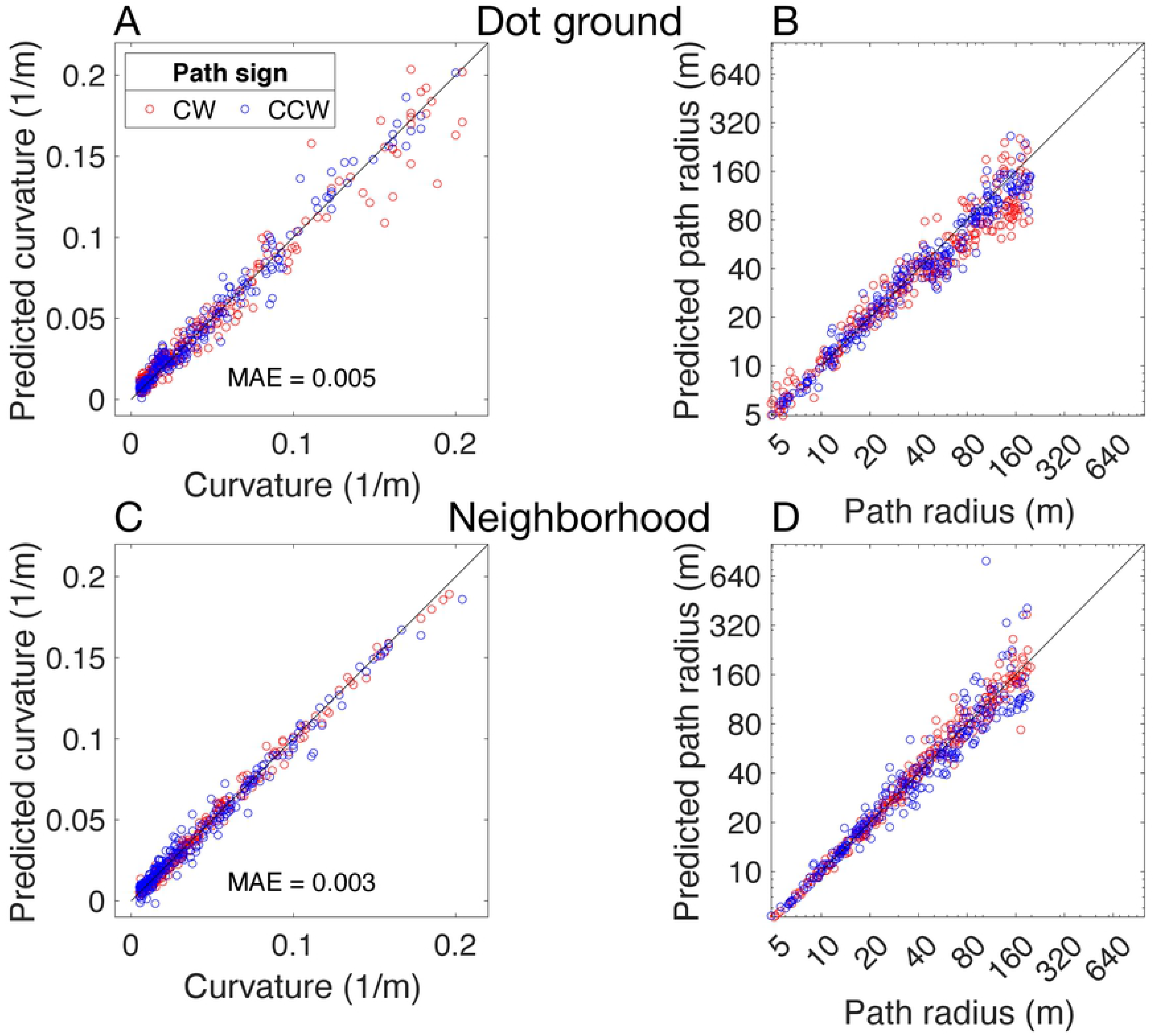
Path curvature estimates decoded from model MSTd in the dot ground (A-B) and neighborhood (C-D) datasets. Top panels depict estimated curvature in the 1/m units used by the deep network decoder for training and prediction. Bottom panels represent the same values as path radius estimates (m units). Estimates for CW curvilinear paths are shown in red and those for CCW paths are shown in blue.

**Figure 5:**
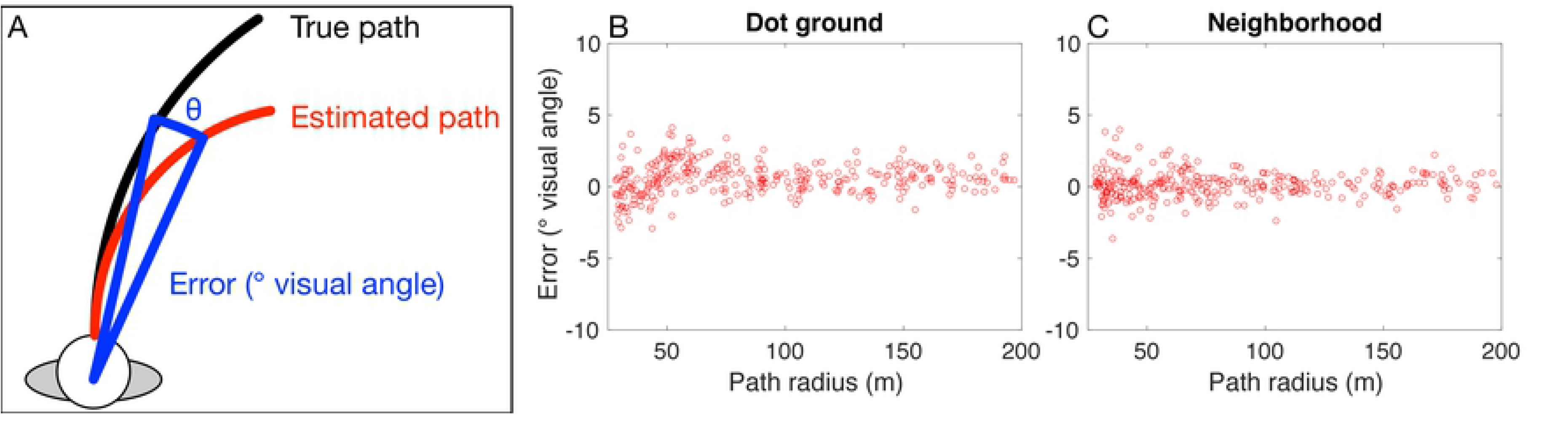
Error in estimated path curvature expressed as degrees of visual angle. (a) Error computed as the angular difference between the true curvilinear path (black) and that estimated by the decoder (red) at a distance of 10 m from the observer. (b) Error in visual angle for the dot ground dataset. Path radii 28-200 m are shown. (c) Same as (b) for the neighborhood dataset.

**Figure 6:**
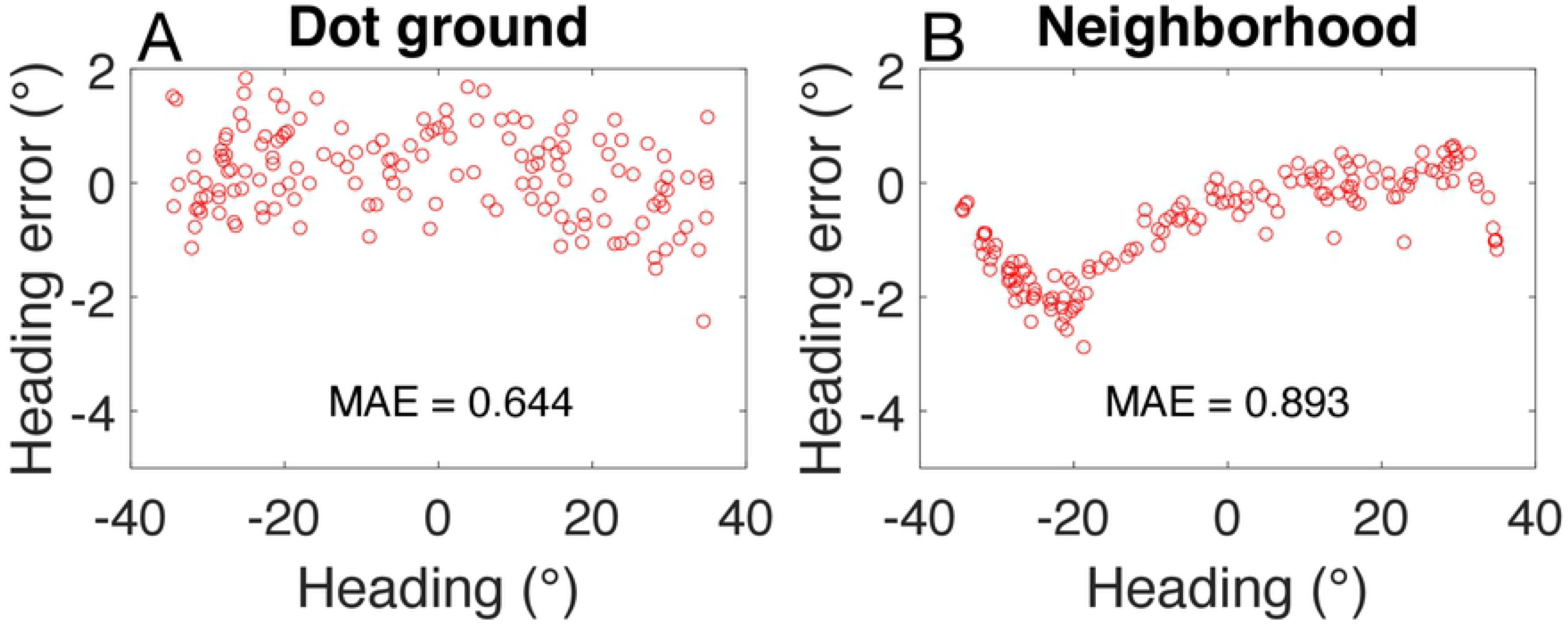
Error in estimated heading in the dot ground (a) and neighborhood (b) datasets.

Figure 3 shows that gaze offset angle was accurately decoded from activation produced by the spiral cell model population in the dot ground (Figure 3a) and neighborhood datasets (Figure 3b). In the dot ground environments, the mean absolute error (MAE) was only 1.5° and estimates generally fell within ±5° of the true value for virtually all optic flow samples. Estimates for CW (red) and CCW (blue) were comparable and error did not depend on the magnitude of the gaze offset. Path sign was decoded from the spiral cells with 100% accuracy.

In Figure 4, we compare path curvature decoded from model MSTd with ground truth values. Path curvature was decoded with a high degree of accuracy in both the dot ground (Figure 4a; MAE = 0.005; R^2^ = 0.99) and neighborhood datasets (Figure 4c; MAE = 0.003; R^2^ = 0.99). It is noteworthy that we trained the decoder to estimate path curvature (1/m) rather than the perhaps more intuitive measure of path radius (m). We made this choice because of the nonlinear relationship radius has with path curvature (Eq. 1):

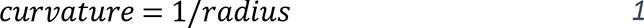

This implies that fixed differences in path radius when the path radius is small (e.g. 4 vs 5 m) have larger effects on the optic flow than when the path radius is large (e.g. 84 vs 85 m) (Figure 1b). We replotted the same data converted to path radii to aid interpretability (Figure 4b,d). Figure 4b reveals increased variability and an underestimation bias for large path radii (small path curvature). This means that in an extreme case, for example, a ∼80 m path was estimated as a ∼50 m path. While this may seem like a large error, there are only subtle differences in the optic flow features that drive the estimates (compare center and right panels of Figure 1b).

To better appreciate the decoded estimates of high radii paths, we computed the difference in visual angle subtended between the true and decoded the paths (Figure 5a). Small angular differences imply close proximity between the two paths from the observer’s vantage point. We measured this difference 10 m from the observer, a distance that has been used to obtain human curvilinear path judgments (Li & Cheng, 2011). Given the focus of this analysis on high radii paths, we followed Li & Cheng and considered paths radii 28 m and greater. Figure 5b-c show that for both datasets the error in visual angle between the true and decoded paths is small and independent of path radius. This means that the estimated path closely tracks the true path, regardless of its radius. That is, decoding precision is actually comparable across path curvatures and is only limited by the lack of distinction in the optic flow inputs for high radii paths.

Next, we considered the accuracy with which the neural system decodes heading during linear self-motion. Note that this is equivalent to the gaze offset angle during curvilinear self-motion, but we use the term “heading” when referring to self-motion along a straight path for consistency with the broader literature. Figure 6 shows errors in heading estimates decoded from model ground cells using the same convolutional deep network decoding architecture used in Branch 1 of the path curvature decoding network (see Methods). Heading accuracy in the dot ground (MAE = 0.644°) and neighborhood scenes (MAE = 0.893°) is comparable to the 1-2° precision of human judgments (Warren, Morris, & Kalish, 1988).

### Temporal stability of self-motion estimates

The decoding networks were calibrated using neural signals present at a fixed time (frame 10) relative to stimulus onset, yet the signals change over time. We sought to characterize the stability of each decoded estimate over the course of the 5 sec (150 frame) videos.

The time series in Figure 7a shows the path curvature decoded from the network on each frame of a video wherein the simulated observer that moves along a 25 m path (0.04 path curvature) and gaze offset of −12.5°. Despite the evolving optic flow field, the decoded path curvature estimate achieves and maintains a steady state of ∼0.055±0.01 (1/m) (16.67 – 25 m) after the first several frames. This ∼7 frame “warm-up” period reflects the time required for the optic flow signal to propagate throughout all layers of the neural model and for dynamics to stabilize for the first time. We demonstrate below that this is a one-time occurrence that only arises at the outset of the simulation that does not impede sensitivity to change at later times.

**Figure 7:**
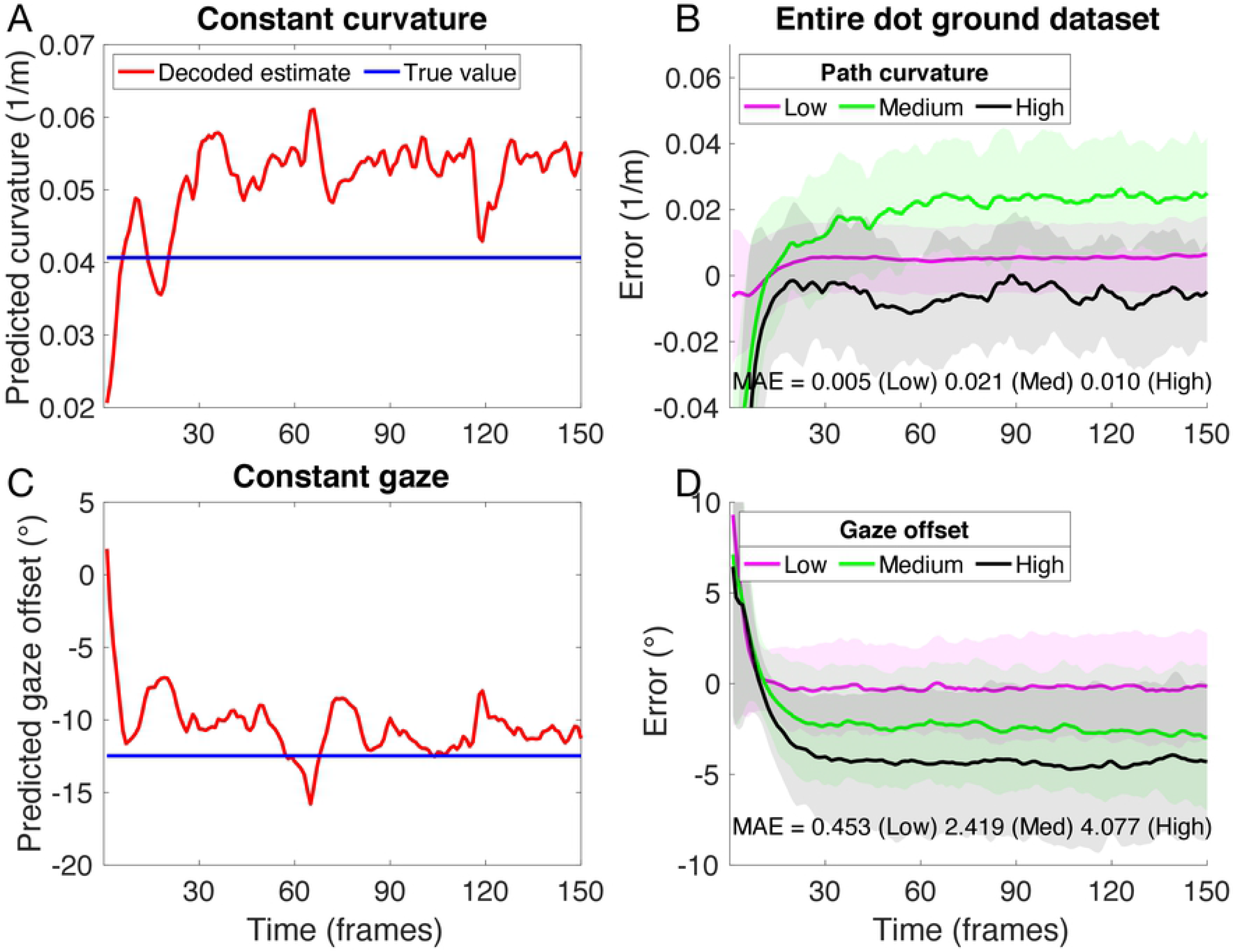
Curvature (A,B) and gaze offset (C,D) predictions decoded from model MSTd over 5 sec durations in the dot ground dataset. Left column: time series of estimated curvature (A; red) and gaze offset angle (C; red) from a single video exemplar. The true path curvature and gaze offset (blue) remained invariant over time. Right column: error time series showing estimated curvature (B) and gaze offset (D) averaged across all samples. Error is computed as predicted value – true value. Colored curves correspond to mean estimates for video samples binned into thirds (low, medium, high) with respect to path curvature and gaze offset. Colored bands correspond to ± 1 standard deviation.

Figure 7b plots the mean error in decoded path curvature across all videos in the dot ground dataset, grouped by dividing the 0.005—0.2 range of tested curvatures into thirds (“low”, “medium”, and “high” curvature bins). The decoded estimates in each curvature range exhibit comparable stability and do not greatly fluctuate over time. It is noteworthy that the deep learning decoders do not explicitly account for temporal continuity across samples — signals decoded at each instant are treated independently in time. Temporal integration and competitive dynamics in MSTd within the CD neural model enhance stability by ensuring that self-motion signals in MSTd generate similar patterns of activation despite fluctuations due to the evolving optic flow field (Layton & Fajen, 2016b).

Figure 7c-d presents the same analysis applied to the gaze offset angle estimated at each frame of video. The decoded values also demonstrated temporal stability and a high degree of accuracy across all samples. Figure 7d shows that larger gaze offset angles produce larger mean errors over time than smaller gaze offset angles, but MAE does not exceed 5°.

More broadly, Figure 7b,d reveals that the ∼7 frame “warm-up” period arises across the dataset. This might suggest that the decoded estimates could be sluggish to adapt to changes in observer’s self-motion over times. We tested this using a “multi-segment” video dataset wherein the simulated self-motion trajectory abruptly transitions between low to high path curvature curvilinear paths every 30 frames within single videos. Gaze offset remained constant throughout each multi-segment video. Figure 8a demonstrates that decoded estimates rapidly adapt to changes in path curvature beyond the initial “warm-up” period, independent of the magnitude of the shift. The decoded CW/CCW path sign is reflected in the sign of the plotted path curvature estimate. In Figure 8b, we compared the decoded gaze offset angle time series with the true value (― 10°). After the initial “warm-up” period, estimates generally fell within ± 5 of the true value. Error in the decoded estimate spiked as the path radius changed from 500 m to 5 m at frame 90, though accuracy of estimates rapidly improved after this transient excursion. Figure 8c,e show the time series of curvature predictions for other multi-segment videos that contain the same transitions among path curvatures, but different gaze offset angles held constant throughout each video — 5° and 0°, respectively. Estimates demonstrate consistency across the different gaze offset conditions. Figure 8d,f show the time series of gaze offset angle predictions corresponding to the same videos, which demonstrate similar accuracy and temporal stability.

**Figure 8:**
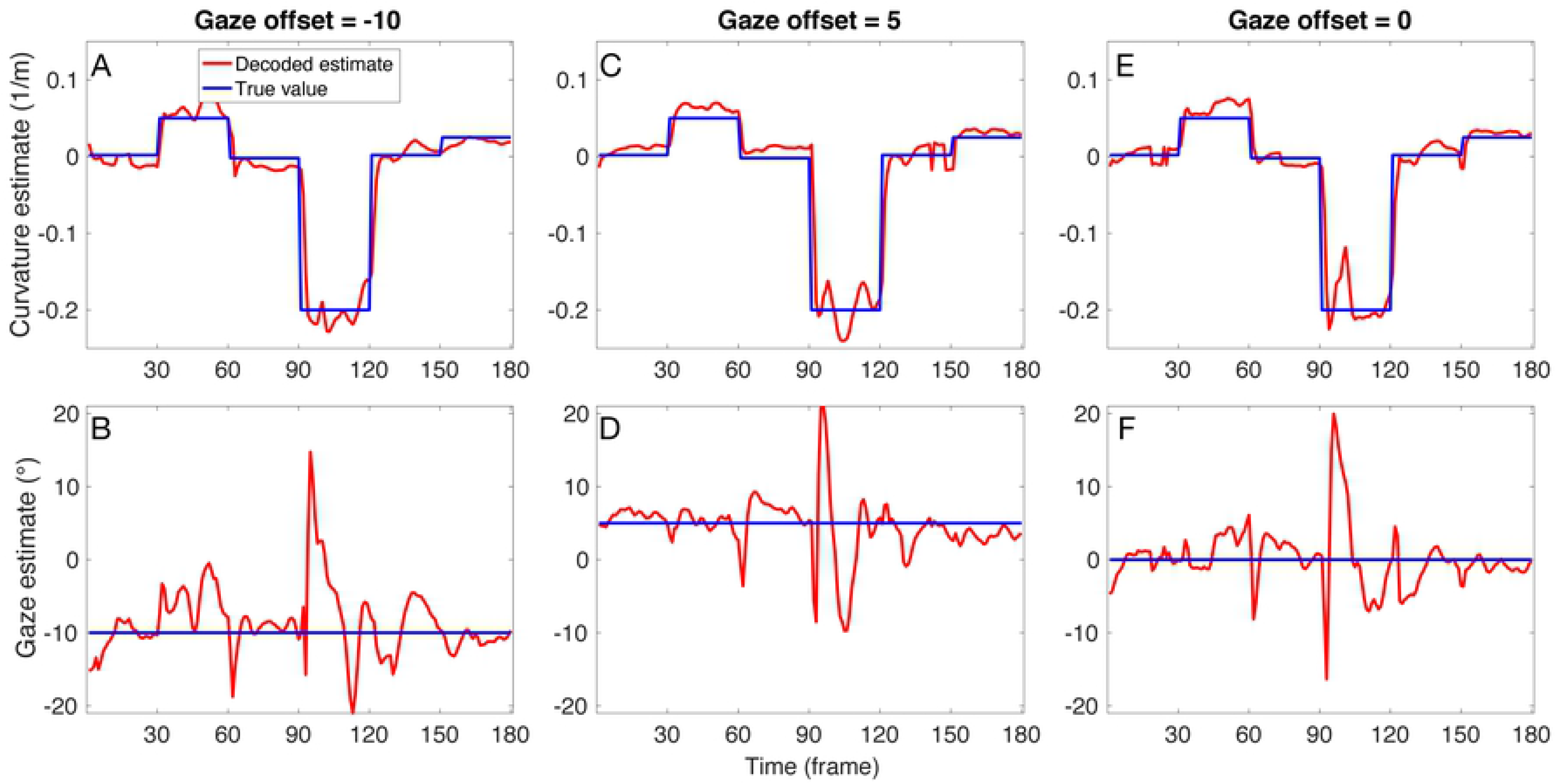
Time series estimates of path curvature (A,C,E) and gaze offset angle (B,D,F) from the multi-segment dataset. Each column corresponds to simulations of the same video, each out which had a consistent gaze offset angle (left: 10°, center: 5°, right: 0°), but path curvature that changed in magnitude and CW (+1) and CCW (−1) sign every 30 frames. Blue curves show the ground truth curvature and gaze offset values. Red curves show values decoded from neural model at each frame.

### Comparison to mathematical model of curvilinear rotation estimation

We compared the performance of our system to another vision-based approach that estimates curvilinear self-motion from optic flow. We selected the model of Perrone (2018) because its parameters were fit using constraints informed by the primate visual system and the precision of human self-motion judgments. It is noteworthy that the Perrone model is a mathematical algorithm rather than a neural model: it applies a mathematical vector projection on each flow vector throughout the visual field to recover the rotational component in the optic flow (see Perrone, 2018 for details). While the model can operate with only optic flow signals, Perrone showed that it must leverage one or more nonvisual signals (e.g. vestibular) to account for human-like self-motion estimation. Because our system presently only works with optic flow, we compared performance to the Perrone model without the nonvisual component (vestibular signal set to 0). We simulated our neural model with optic flow generated according to specifications of the “Vision Only” Experiment 3 of Perrone (2018), with the exception that we used a ground plane rather than a 3D dot cloud for consistency with our prior simulations. See Methods for a full description of the neural decoder training and testing protocol used for this dataset.

We parameterized model estimates with respect to rotation rate in the optic flow caused by the observer’s curvilinear self-motion. Rotation rate (*R*) is the metric used by Perrone (2018) and can be computed using the path curvature, which was estimated by the decoder, and the observer’s self-motion speed (*T* = 1.5 *m*/*sec*) as follows:

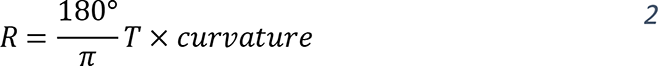

The first row of Figure 9 emulates the format of Figure 13a-b of Perrone (2018) and shows path rotation estimates (Figure 9a) alongside the distribution of errors (Figure 9b) decoded from model MSTd. We obtained a root-mean-squared (RMS) error of 0.881°/sec on the test set, which is comparable to that found by Perrone (RMS = 0.91°/sec).

**Figure 9:**
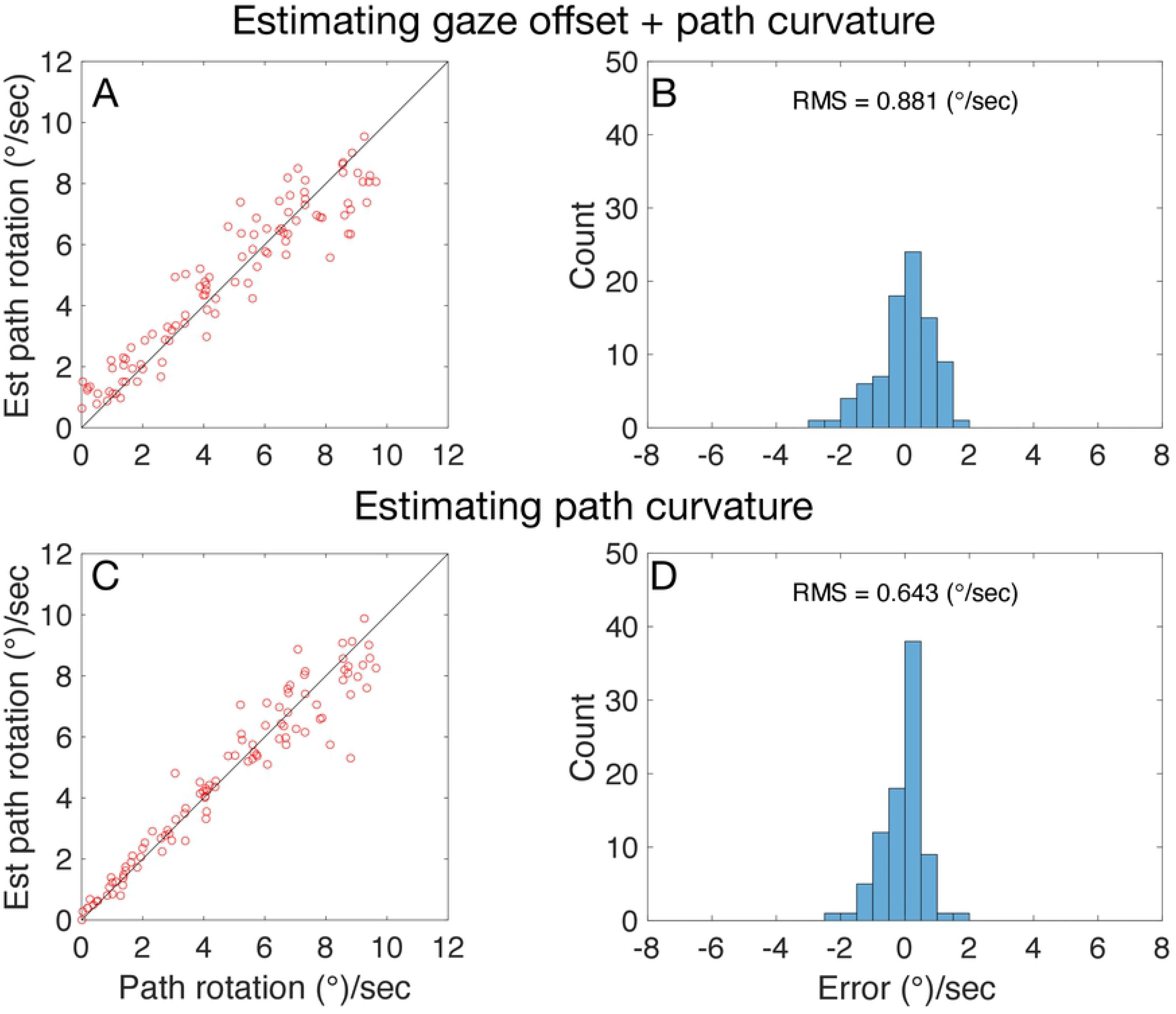
Decoded estimates of curvilinear path curvature expressed as path rotation rate (°/sec) experienced by observer moving at a speed of 1.5 m/sec along CW paths. Top (a,b): Performance when decoder estimates both gaze offset and path curvature. Bottom (c,d): Performance when decoder only estimating path curvature. Histograms in right column show errors corresponding to values shown in left column. RMS represents root-mean-squared error.

In these simulations our system estimates both gaze offset and path rotation (via path curvature), whereas the corresponding simulations of Perrone (2018) focuses only on the latter. When we configure the decoder to estimate path rotation alone by supplying the ground-truth gaze offset angles (Figure 9c-d), RMS error decreases to 0.643°/sec. This comes approximately halfway in closing the gap in performance between the vision-only (RMS = 0.91°/sec). and joint vision-vestibular model of Perrone (RMS ≈ 0.3°/sec).

### Comparison to human judgments

We also compared model path curvature estimates to human judgments of curvilinear self-motion. Our aim was not to perform a thorough analysis of the existing psychophysical literature on curvilinear self-motion perception; rather we sought to compare a subset of conditions already tested to human data obtained under similar circumstances. We simulated the “gaze-along-heading” condition from Li & Cheng (2011) because it approximates a subset of conditions from the dot ground dataset: simulated CW and CCW curvilinear self-motion over a ground plane of dots with a gaze offset of 0°. Accordingly, we used the decoder trained on the dot ground dataset to estimate path curvature in the gaze-along-heading condition (see Methods).

Figure 10 compares model path curvature estimates with judgments made by human subjects in Li & Cheng (2011). We expressed decoding error in terms of the previously used visual angle error measure to afford direct comparison (Figure 5). Consistent with Li & Cheng (2011), positive error indicates an overestimation of the path curvature. Figure 10 demonstrates that the model produces human-like estimates when gaze is aligned with the instantaneous heading direction (gaze offset of 0°). Negative model error in the high path curvature condition reflects the tendency of the decoder to slightly underestimate path curvature within that range of path curvatures. Overall, these simulations demonstrate that path curvature estimations decoded from model MSTd can exhibit comparable accuracy to humans.

**Figure 10:**
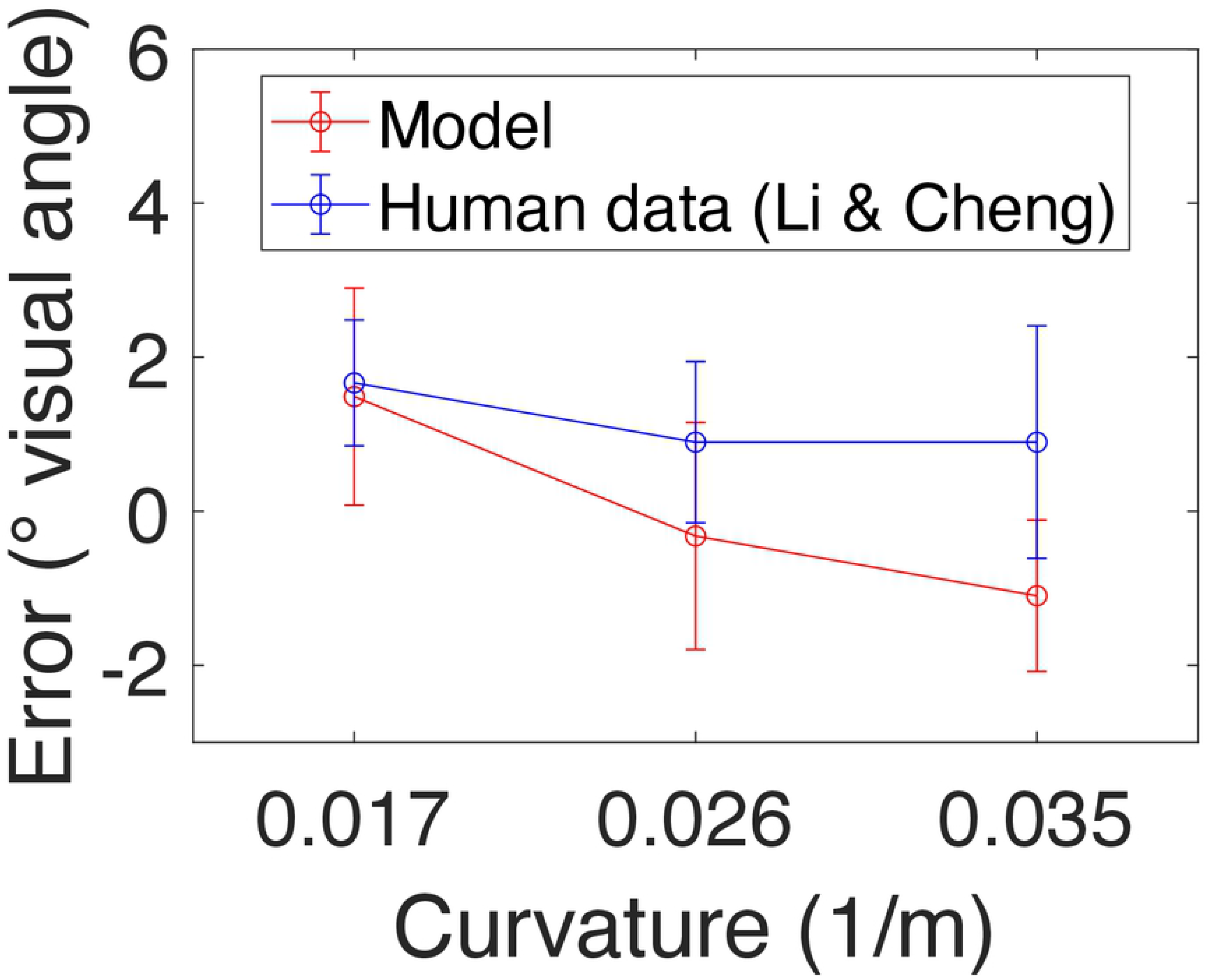
Model path curvature estimates compared with human judgments (Li & Cheng, 2011). We simulated the gaze-along-heading condition of Li & Cheng simulating curvilinear self-motion over a ground plane of dots with a gaze offset of 0°. Error expressed as difference in degrees of visual angle between the estimated and actual paths 10 m from the observer (estimated – actual; same as Figure 5). Error bars on model estimates indicate ±1 standard deviation while those on human judgments indicate ±1 standard error.

## Discussion

Neurophysiology of MSTd has focused on heading during linear self-motion and cells tuned to radial optic flow; tuning to spirals and other complex motion patterns have long been known (Saito et al., 1986) (Tanaka, Sugita, Moriya, & Saito, 1993) (Graziano et al., 1994), but their function have been less clear. The aim of the present study was to investigate whether populations of units that model the heterogenous pattern tuning in MSTd could estimate a broader range of parameters that describe the observer’s self-motion. We have demonstrated accurate estimation of the following self-motion characteristics during linear and curvilinear self-motion when using deep learning to decode signals produced by a mechanistic neural model of MSTd:

- Heading from cells tuned to ground flow.
- Gaze offset angle, CW/CCW curvilinear path sign, and path curvature from distributed activation produced from cells tuned to diverse patterns, including full-field and hemi-field spirals, ground flow, and radial expansion.

Estimates were accurate and stable when decoded across time from synthetic and more realistic scenes.

### Linear and curvilinear self-motion signals in MSTd

We decoded heading from MSTd cells tuned to different FoE positions following strong connections forged over decades of physiological (Duffy & Wurtz, 1995) (Gu et al., 2012) (Gu et al., 2006) (Britten & van Wezel, 2002) and computational (Warren & Saunders, 1995) (Royden, 2002) (Layton et al., 2012) studies. These studies tend to map MSTd RFs using full-field radial expansion that resemble optic flow that an observer would experience during linear self-motion toward fronto-parallel planes or a 3D dot cloud. We derived estimates from model units tuned to ground flow rather than radial expansion due to their appropriateness for the naturalistic ground-based self-motion employed in the datasets. Tuning in monkey MSTd to optic flow generated from self-motion over a ground plane has been previously demonstrated (Logan & Duffy, 2006). Decoding from cells tuned to radial expansion should yield qualitatively similar, albeit perhaps less precise, results given that ground flow yields weaker and less selective radial cell activation.

Simulations of the CD neural model informed our selection of MSTd cell populations from which to decode the other self-motion parameters. The use of ground tuned MSTd cells to decode path curvature emerged from simulations demonstrating systematic bias proportional to path curvature (Figure S1c). This dependance has been previously noted, but not thoroughly investigated (Cameron, Grossberg, & Guenther, 1998). While Graziano and colleagues have hypothesized that spiral cells may play a role in estimating self-motion or object motion as the observer fixates a stationary point in the environment (Graziano et al., 1994), we are not aware of an existing proposal linking the estimation of gaze offset angle or path sign during curvilinear self-motion.

In contrast to the dependence on both spiral and radial (or ground) cells in our decoding network for path curvature, the neural model of Layton & Browning (2014) relies only spiral cells to explain human curvilinear path judgments. There are several key differences between our models and studies. First, Layton & Browning only considered a limited range of path radii (28—58 m) to facilitate comparison with human psychophysical data. It is possible that spiral cells alone could estimate with human-like accuracy within this range. Second, the Layton & Browning model does not explicitly account for gaze offset so joint variations in path curvature and gaze offset would likely confound the model. Third, Layton & Browning did not obtain estimates of path curvature and gaze offset in metric units as we did here. Instead, the model mapped internal differences in MSTd unit activation along a spiral continuum that resembles Figure S1b onto errors in path judgments made by humans. To do this, they anchored model performance to MSTd activity obtained for each path radius when gaze was tangent to the future path (“gaze-along-heading”; 0° gaze offset), when humans are accurate in their judgments. That is, the model computes error for a given path radius by subtracting the position of the activity peak in spiral space from that produced in the gaze-along-heading condition. This simplistic approach also does not actually estimate path curvature — for example, it cannot indicate that a 5 m path is a 5 m path or differentiate a 5 m path from a 25 path. The comparisons that the Layton & Browning model make relative to a known baseline path curvature are much less demanding than the precise estimates obtained in the present work.

### Decoding signals with deep learning

To better appreciate the role of each cell population in estimating respective self-motion parameters, we designed an ensemble of specialized deep learning decoders rather than a single decoding network. This approach helps expose the contributions of each cell population and mitigates some issues surrounding the interpretability of representations learned by deep learning networks (Serre, 2019) (Grossberg, 2020). Hierarchical dependencies among the decoders increases the specificity of our hypotheses about the function of different MSTd cell subpopulations. In particular, the decoder network making the gaze offset predictions depends on the predictions made by the path network. Moreover, the path curvature network depends on estimates produced by both the path sign and gaze offset networks (see Methods). Our use of the ensemble of decoder networks, each specialized for estimating a single parameter, critically depended on the specificity of hypotheses about the role of MSTd subpopulations.

That is, without MSTd signals from units with specific tuning characteristics and a principled way to cascade the predictions produced by each decoder network, we would have required a large, more monolithic decoding network that relied on pixel or optic flow inputs. The MSTd signals generated by the CD neural model could be seen as intermediate “features”, which confer benefits for training deep networks, such as decreased training time (Zhou, Krähenbühl, & Koltun, 2019) and relatively few numbers of layers (7-9) and number of units per layer (20-40; see Methods) by modern standards (Alom et al., 2018) (Krizhevsky, Sutskever, & Hinton, 2012) (Simonyan & Zisserman, 2014).

### Limitations

Our study demonstrates the sufficiency of decoding linear and curvilinear self-motion parameters from the hypothesized MSTd subpopulations, it does not show that the proposed associations are necessary for accurate estimation. For example, decoding from other combinations of MSTd subpopulations could possibly yield similar or even more accurate results. Exploring the large number of configurations of decoding networks, approaches to decoding, and combinations of MSTd subpopulations that could support comparably accurate estimates falls outside the scope of the present investigation. This analysis is the focus of a forthcoming study that complements the present work.

The feedforward convolutional deep network architecture used in the present study offers state-of-the-art performance when decoding neural signals from many cortical areas (Glaser et al., 2020) that is superior to traditional approaches, such as Wiener filters, Naïve Bayes, support vector machines, and Kalman filters. Despite the performance benefits, deep learning, however, does not clearly expose the mapping that is learned between neural activity and the desired parameter value. The lack of a specific closed-form mathematical mapping makes it challenging to ascertain the contributions of different MSTd populations when estimating self-motion parameters. This contrasts with simple decoding strategies, such as winner-take-all (Layton et al., 2012) (Royden, 2002) and population vector (Layton & Fajen, 2020) that propose straightforward mappings between heading estimates and radial cell activation: cells tuned to the heading of the observer produce larger activation than those tuned to dissimilar headings.

The present study does not address the influence of eye movements, which have been shown to modulate MSTd signals during self-motion (Danz et al., 2020) (Shenoy et al., 1999) (Inaba, Shinomoto, Yamane, Takemura, & Kawano, 2007) and human self-motion judgments (Royden, Crowell, & Banks, 1994) (Warren & Hannon, 1990) (Cheng & Li, 2012). Eye movements introduce an endogenous source of rotation into the optic flow field that is independent from the rotation caused by curvilinear self-motion. To estimate curvilinear self-motion during eye movements, the prevalent efference copy theory and related models propose that the visual system relies on nonvisual signals (e.g. vestibular signals specifying eye position) to factor out the rotation due to eye movements, leaving on the rotation due curvilinear self-motion (Perrone, 2018) (Perrone & Krauzlis, 2008). Such a mechanism could be used to remove the effects of rotation prior to decoding from our model. However, the extent to which MSTd decomposes the optic flow field into distinct translational and rotational components visually (Orban et al., 1992) or in the presence of vestibular signals (Danz et al., 2020) has been challenged. The mechanism by which the visual system tolerates rotation in optic flow during self-motion remains unclear.

Some have examined linear heading perception when gaze remains fixed and eye movements are simulated within the virtual environment of the display (“simulated eye movement condition”) (Royden et al., 1994). In the absence of vestibular signals that accompany real eye movements, humans appear to treat the simulated rotation as curvilinear self-motion without eye movements (Banks, Ehrlich, Backus, & Crowell, 1996). This suggests that rotation due to curvilinear self-motion is not factored out, decomposed, or removed, at least for the purposes of estimating curvilinear self-motion. The system presented here is consistent with this proposal, since neither the neural model nor decoding networks explicitly remove rotation due to curvilinear self-motion. Indeed, pattern tuning in MSTd may emerge through the dimensionality reduction that neurons perform on their motion inputs (Beyeler, Dutt, & Krichmar, 2016) rather than a process whose purpose is to segment translation from rotation.

### Conclusion

We have presented a system that accurately estimates the parameters describing the observer’s self-motion along linear and curvilinear paths from a neural model that processes optic flow. Estimates emerge by decoding signals from units that emulate the diverse motion pattern tuning of primate MSTd. The system produces accurate and stable estimates in random-dot textured and more visually realistic scenes over time that adapt to changes to the observer’s self-motion. Our findings support the possibility that MSTd could jointly signal the observer’s linear and curvilinear self-motion.

## Methods

### Optic flow datasets

Table 1 summarizes the optic flow datasets that consisted of 30 frame-per-second digital videos of linear and curvilinear self-motion through random dot and photorealistic virtual environments at 512×512 resolution. For datasets focusing on curvilinear self-motion, the observer moved along a circular path. We equated the parameters specifying self-motion across the dot ground and neighborhood environments to facilitate comparison.

**Table 1:**
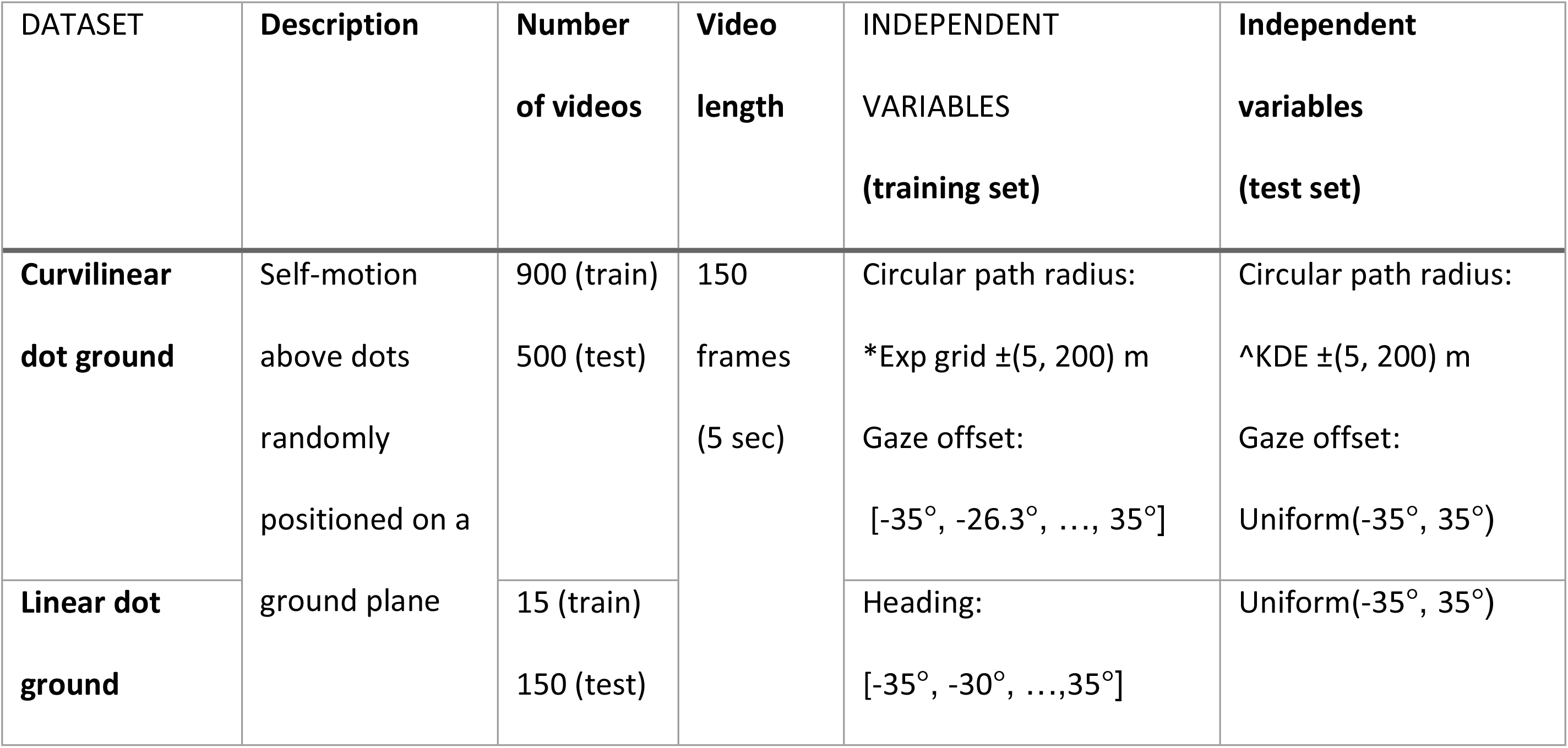
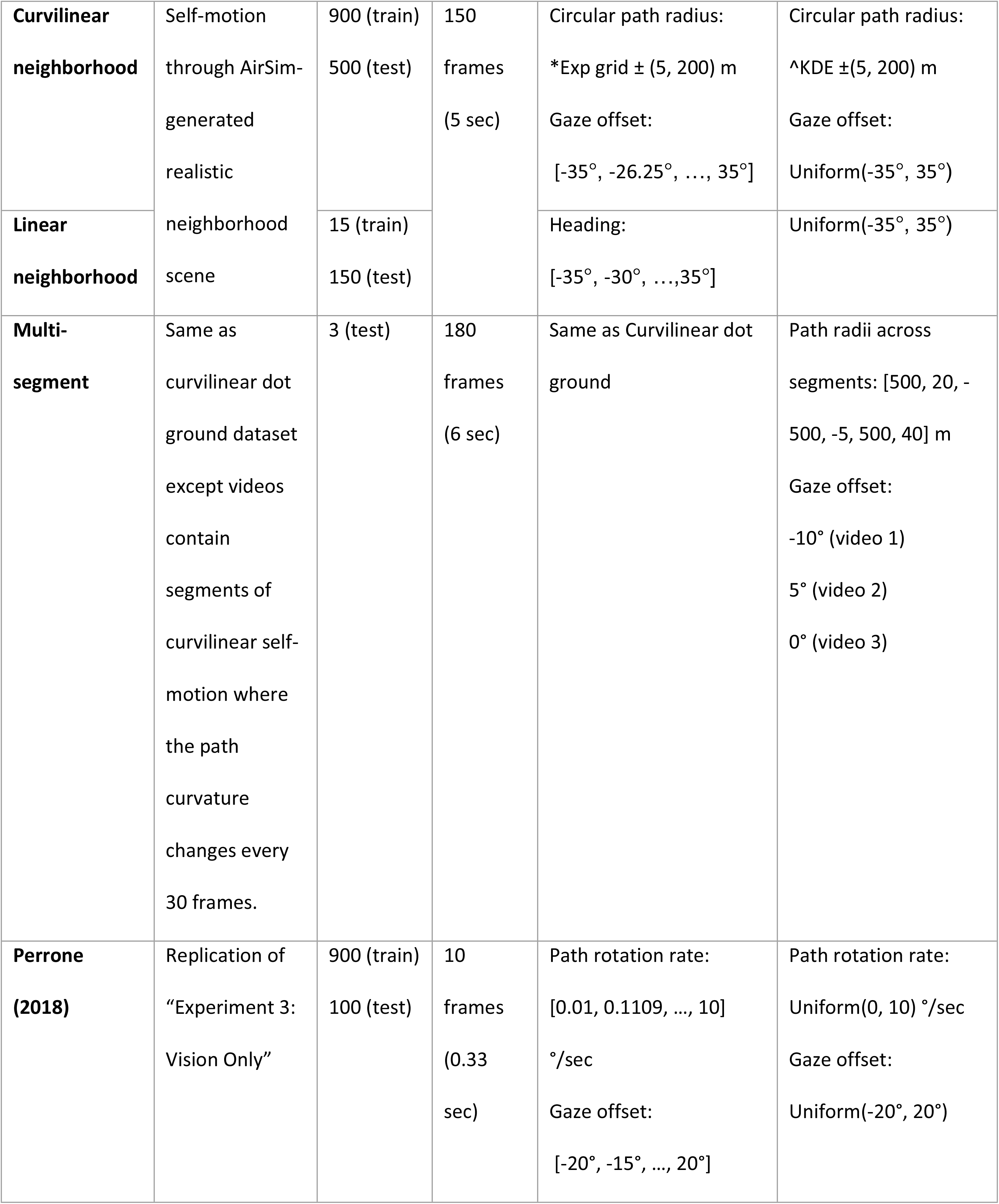
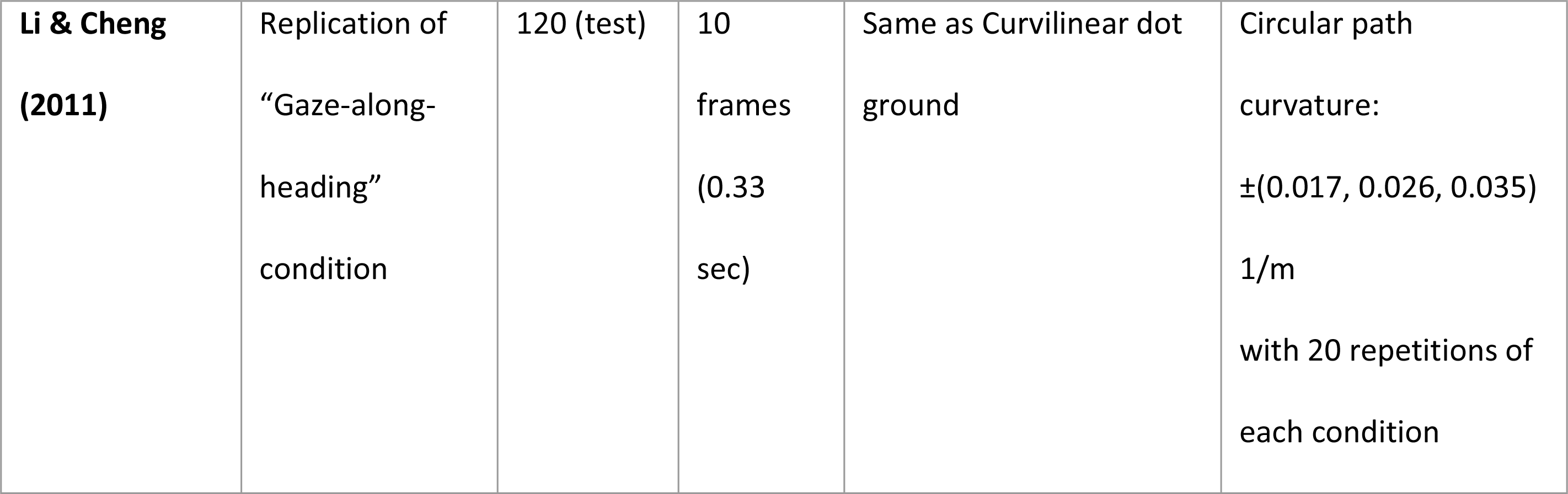
Description of optic flow datasets used in simulations. Negative gaze angles indicate gaze offset to the left of the straight-ahead. Negative path radii indicate curvilinear self-motion toward the left, along a CCW path. *Exp grid indicates that values are regularly spaced in exponential space between the specified endpoint values. ^KDE indicates that values were sampled via kernel density estimation using an empirical probability distribution based on the training set values (see main text for details). Uniform indicates that values were sampled from a uniform distribution between the specified endpoints. Unless indicated otherwise, we decoded from frame 10 of the dot ground and neighborhood datasets.

### Dot ground datasets

On each video frame in the dot ground datasets, we computed the optic flow using a pinhole camera model (Raudies & Neumann, 2012) and standard analytic equations (Longuet-Higgins & Prazdny, 1980). Table 2 summarizes the parameters that specify the simulated observer and environment in the analytic ground dataset. We clipped and replaced dots that exited the field of view or valid depth range to ensure that the same number of dots always remained visible. Figure 2b shows example optic flow fields from the dot ground datasets.

**Table 2:** Parameters specifying self-motion through dot ground environment

| Parameter | Value |
| --- | --- |
| Spatial resolution | 512 x 512 pixels |
| Translational speed | 3 m/sec |
| Focal length | 1.74 cm |
| Field of view | 90° |
| Eye height | 1.61 m |
| Dots on ground plane | 2000 |
| Observer-relative depth range of dots<br>(min, max) | (1 m, 50 m) |

### Neighborhood datasets

We generated the neighborhood datasets using Microsoft AirSim, a simulation environment for drones that renders photorealistic scenes using the Unreal game engine (Shah, Dey, Lovett, & Kapoor, 2017). We used a rich outdoor neighborhood scene with grass, trees, houses, fences, and streets, and other objects (Figure 2c). After generating videos with self-motion parameters that matched those used in the dot ground datasets, we computed optic flow using DeepFlow2 (Weinzaepfel, Revaud, Harchaoui, & Schmid, 2013).

### Training sets

In the training sets we sampled self-motion parameters (gaze offset, heading) on a regular grid to ensure balanced coverage. We sampled path radius differently than the other parameters due to the nonlinear relationship between circular path radius and curvature (Eq. 1). We accounted for this by sampling small radii more finely than large radii using the following recursive generating formula:

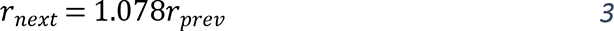

We used a starting radius of 5 m and selected the coefficient so that Eq. 3 would generate 50 radii between 5–200 m. We doubled the number of radii by including both CW and CCW versions of each path.

### Test sets

Gaze offset and heading were randomly sampled from a uniform distribution in the test sets (Table 1). We used kernel density estimation (KDE) to randomly sampled path radii in proportion to the training set values. As in the training set, this biased values toward smaller radii that produce larger variation in path curvature. We fit the path radii used in the training set to an empirical probability density function via the MATLAB function *ksdensity*, which we then sampled within the training set range with 0.1 m granularity. As in the training set, we also included CCW versions of these paths.

### Neural model of MT and MSTd

We processed the optic flow datasets with the Competitive Dynamics neural model of MT and MSTd (Figure 11) (Layton et al., 2012) (Layton & Fajen, 2016b) (Layton & Fajen, 2020) (Layton & Niehorster, 2019). Populations of model neurons that emulate a range of physiological properties transform the optic flow into neural signals from which we subsequently decode the parameters describing the observer’s self-motion. The CD model accounts for human judgments of self-motion, including in the presence of independently moving objects (Layton & Fajen, 2017) (Layton & Fajen, 2016b), and object motion during self-motion (Layton & Fajen, 2020) (Layton & Fajen, 2016a) (Layton & Niehorster, 2019). We made several key changes to the most recent version of the model (Layton & Niehorster, 2019) (Layton & Fajen, 2020):

- Given focus in the present work on self-motion estimation, we only simulated the requisite MT^+^–MSTd pathway of the model, excluding the MT^-^–MSTv object motion pathway (Layton & Niehorster, 2019) (Layton & Fajen, 2020).
- We simulated units tuned to direction and speed only, omitting disparity tuning.
- MT^+^ neurons tuned to faster speeds possessed larger receptive fields (RFs) than those tuned to slower speeds (Mikami, Newsome, & Wurtz, 1986) (Tanaka et al., 1986).
- Speed tuning in MT adapted based on the recent time history of the optic flow signal rather than remaining fixed (Steinmetz, Layton, Powell, & Fajen, 2021).
- Each MSTd unit received input from a random subset of the MT^+^ units. In prior versions of the model, MSTd units were fully connected to the constellation of MT^+^ units that compose the preferred pattern.
- To better manage the greater diversity of cell types simulated (e.g. radial, spiral), we divided model MSTd into several processing layers. Within each, we grouped units according to their optic flow pattern preference (Figure S1).

**Figure 11:**
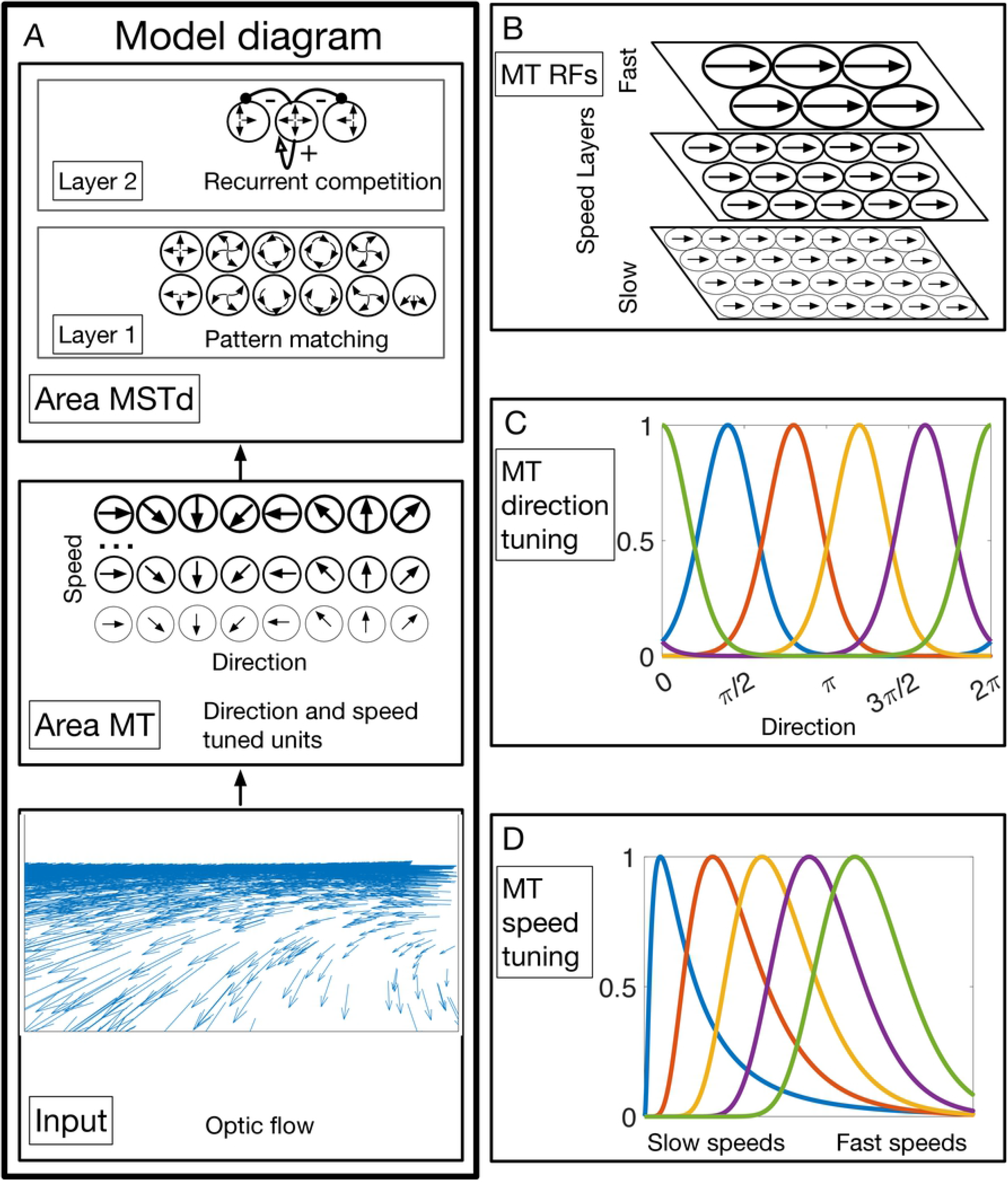
Overview of the Competitive Dynamics neural model of MT-MSTd. (A) Model area MT contains direction and speed tuned units. Units in model area MSTd integrate motion signals from MT consistent with the preferred motion pattern and compete with one another. MSTd units are tuned to full and hemi-field motion in a pattern continuum: radial, CCW spirals, CCW centers, CW spirals, CW centers, and ground flow. (B) MT units tuned to faster speeds possess larger RFs and process the visual field at lower spatial resolution than those tuned to slower speeds. (C) Example MT direction tuning curves. (D) Example MT speed tuning curves. They adapted through dynamic efficient sensory encoding to optimize responsiveness over the changing dynamic range of input signals over time.

These changes and the complete model details will be fully described in the following sections. Figure 11a summarizes the architecture of the CD model simulated here, Table 3 specifies the values of parameters used to simulate model MT, and Table 4 specifies those used to simulate model MSTd.

**Table 3:** Parameters used in model area MT.

| Parameter | Description | Value |
| --- | --- | --- |
| $\vec{s}^{MT}$ | Optic flow spatial scaling factors for each speed layer.<br>Lower indices correspond to cells tuned to slower speeds. | [8, 8, 16, 16, 16]<br>Corresponding spatial resolution:<br>$s_{MT} = 8$: 64x64<br>$s_{MT} = 16$: 32x32 |
| $N^s$ | Number of preferred speeds | 5 |
| $p^{MT}$ | Gaussian pyramid kernel diameter-to-standard deviation<br>scale factor | 1/4 |
| $N^d$ | Number of preferred directions | 24 |
| — | Shape of MT<sup>+</sup> speed layers | $s_{MT} = 8$: 64x64x24<br>$s_{MT} = 16$: 32x32x24<br>Total units across all layers: 270336 |
| $\eta$ | Von Mises direction tuning curve dispersion | 4° |
| $\alpha^{MT}$ | MT<sup>+</sup> passive decay rate | 1 |
| $\tau^{MT}$ | Depressing synapse time constant | 0.1 |
| $\kappa^{MT}$ | Rate at which the synaptic efficacy decreases | 10 |

**Table 4:** Parameters used in model area MSTd.

| PARAMETER | Description | Value |
| --- | --- | --- |
| $N^{MSTd,com}$ | Number of units tuned to CoM positions for each motion pattern (radial, spiral, etc.) arranged in regular across the visual field. | 16x16 = 256 |
| $\vec{\lambda}^{spl}$ | Spirality of patterns to which spiral cells are tuned.<br>Negative (positive) values produce CCW (CW) spirals. | $\pm [0.0, 0.05, ..., 1.0]$ |
| $N^{spl}$ | Number of CW and CCW spiral patterns to which units are tuned. | 21 (CW) + 21 (CCW) = 42 |
| — | Indices of units in pattern tuning continuum. Units corresponding to the endpoints exhibit tuning to radial expansion or CW/CCW center patterns. | CCW full-field spiral (decreasing spirality): 1—21<br>CW full-field spiral (increasing spirality): 22—42<br>CW hemi-field spiral (decreasing spirality): 43—63<br>CCW hemi-field spiral (increasing spirality): 64—84 |
| — | Shape of each MSTd layer | 16x16x85<br>1—84: spiral continuum<br>85: units tuned to ground flow |
| $N^T$ | Number of MT<sup>+</sup> connections within each MSTd pattern template using multivariate Gaussian centered on the CoM. | 200 |
| $\Sigma^{MSTd,T}$ | Covariance matrix of multivariate Gaussian used to sample MT<sup>+</sup> connections within each template. | $s_{MT} = 8$ (MT<sup>+</sup> shape: 64x64x24):<br>$\begin{bmatrix} 1000 & 0 \\ 0 & 1000 \end{bmatrix}$ $s_{MT} = 16$: (MT<sup>+</sup> shape: 32x32x24):<br>$\begin{bmatrix} 500 & 0 \\ 0 & 500 \end{bmatrix}$ |
| $b^{MSTd}$ | Rate at which MT<sup>+</sup> connection weights decrease with distance from the preferred CoM position. | 1e-3 |
| $\alpha^{MSTd,1a}$ | Layer 1a unit passive decay rate | 1 |
| $\vec{\sigma}^{MSTd,1b,com}$ | Layer 1b Gaussian 2D spatial smoothing kernel horizontal and vertical standard deviations | [5, 5] |
| $r^{MSTd,1b,com}$ | Diameter of Layer 1b Gaussian 2D spatial smoothing kernel | 8 |
| $\sigma^{MSTd,1b,pat}$ | Layer 1b Gaussian 1D pattern smoothing standard deviation | 1.5 |
| $r^{MSTd,1b,pat}$ | Diameter of Layer 1b Gaussian 1D pattern smoothing kernel | 7 |
| $\alpha^{MSTd,1b}$ | Layer 1b unit passive decay rate | 1 |
| $\alpha^{MSTd,c}$ | Layer 2 net input adaptive threshold passive decay rate | 1 |
| $\alpha^{MSTd,2}$ | Layer 2 unit passive decay rate | 10 |
| $\beta^{MSTd,2}$ | Layer 2 unit excitatory upper bound | 3 |

### Model area MT

We simulated the population of direction and speed tuned neurons in MT with excitatory surrounds (MT^+^), which project to MSTd (Maunsell & van Essen, 1983) (Berezovskii & Born, 2000) (Born & Tootell, 1992) and influence self-motion signals therein (Yu, Hou, Spillmann, & Gu, 2018).

### MT receptive fields

MT^+^ unit receptive fields (RFs) tuned to like speeds and directions were arranged in a regular spatial grid across the visual field. Units tuned to different speeds occupied different grids. Units tuned to faster speeds possessed larger RFs and were spaced farther apart than those tuned to slower speeds with smaller RFs (Figure 11b). To implement differing RF sizes, we created a Gaussian pyramid based on the 512×512×2 optic flow input, treating the horizontal and vertical optic flow components as two separate images. The image pyramid downsampled the 512×512 images by scaling factors 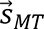 to appropriately match the size of each grid of speed cells. For example, a scaling factor of 8 reduces the original 512×512 spatial resolution to match the 64*x*64 = 4096 grid of units tuned to slow speeds. The vector 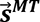 has length 5 (*N*^*s*^) and matches the number of simulated preferred speeds across model MT^+^. We computed the diameter *D*^*MT*^ of the Gaussian pyramid kernel for speed layer *i* according to the following function:

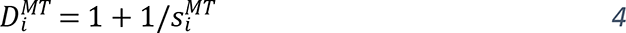

We scaled each kernel standard deviation 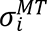 based on the RF diameter:

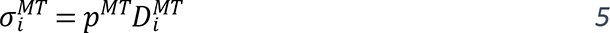

### MT direction tuning

We simulated units tuned to 24 (*N*^*d*^) motion directions (Figure 11c). We modeled each unit’s direction tuning width using the following von Mises distribution

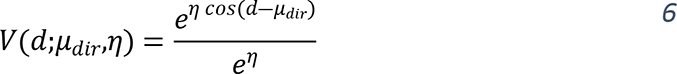

where *d* represents the direction of the optic flow in the receptive field, *μ*_*dir*_ represents the unit’s preferred direction, the shape parameter *η* controls the direction tuning width. We computed the direction of the optic flow vector 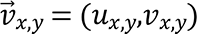 at position (x, y) via the two-argument form of the arctangent function:

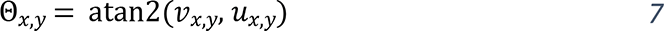

### MT speed tuning

We simulated units tuned to 5 (*N*^*s*^) preferred speeds. Instead of constraining each preferred speed to remain fixed for the duration of each simulation, they adapted to the recent history of speeds present in the optic flow signal. We computed optic flow speed as:

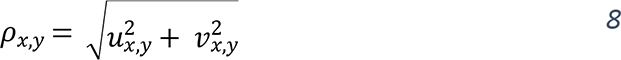

The preferred speeds and their respective tuning curves changed over time to maximize sensitivity to the evolving distribution of speeds present in the optic flow signal (Figure 11d). For example, the 5 speed tuning curves would shift their sensitivity toward faster speeds when the observer nears a tree and redistribute sensitivity toward slower speeds once the tree has been passed. The tuning curves implement a version of efficient sensory encoding (Ganguli & Simoncelli, 2014) that adapts over time. The properties of this dynamic efficient sensory encoding and their impact on self-motion signals are the focus of separate study (Steinmetz et al., 2021).

### MT unit activation

The net input to the MT^+^ unit tuned to speed *s*, direction *d*, and has its RF center at position (*x*, *y*) is the product of the directional *V*(Θ_*x*,*y*_;*d*,*η*) and speed S(*ρ*_*x*,*y*_;*s*) tuning curve outputs:

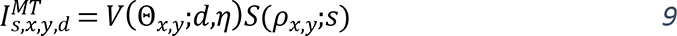

We modeled the dynamics of MT^+^ unit *m*_*s*,*x*,*y*,*d*_ as a simple leaky integrator:

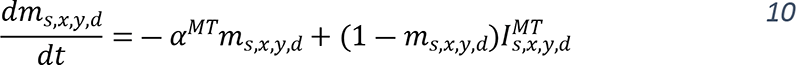

where *α*^*MT*^ is the passive decay rate of the cell.

### MT output

Among models that explain MSTd responses based on their feedforward input from MT, those that include a nonlinearity that compresses MT signals perform best (Mineault, Khawaja, Butts, & Pack, 2012). The compressive nonlinearity could be explained by synaptic depression, the tendency for the same inputs to lose their efficacy over time. We modeled MT^+^ synaptic depression ℎ_*s*,*x*,*y*,*d*_ as follows:

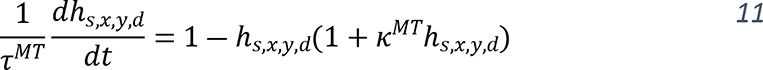

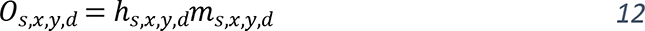

where *O*_*s*,*x*,*y*,*d*_ denotes the output signal from MT^+^ units to MSTd, *τ*^*MT*^ is the synaptic time constant, and *κ*^*MT*^ represents the rate at which the efficacy of the input signal *m*_*s*,*x*,*y*,*d*_ declines over time (Grossberg, 1980).

### MSTd net input

We simulated MSTd cells tuned to radial expansion 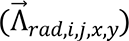 and spiral 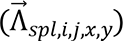optic flow patterns that have been used to characterize MSTd selectivity in neurophysiological studies (Graziano et al., 1994) (Duffy & Wurtz, 1995).

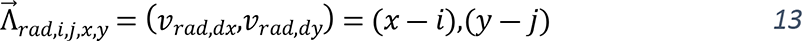

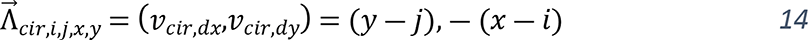

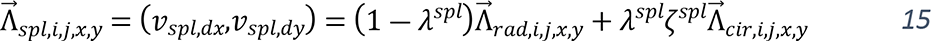

In Eqs. 13-15, (*i*, *j*) indicates the location of the center of motion (CoM), which corresponds to the FoE in the case of radial expansion; (*x*,*y*) indicates visuotopic position of the local motion vector, *ζ*^*spl*^ is 1 for CW spirals and −1 for CCW spirals; and *λ*^*spl*^ is a scalar selected in the range 0–1 that represents the spirality of the pattern. Setting *λ*^*spl*^ = 0 produces a radial expansion flow field, *λ*^*spl*^ = 1 produces a circular center flow field 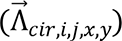, and 0 < *λ*^*spl*^ < 1 creates spiral patterns spanning a continuum between these two extremes. We created MSTd units tuned to *N*^*spl*^ spiral patterns by sampling regularly spaced values of *λ*^*spl*^ (Figure S1a). We simulated MSTd units tuned to both full-field and lower the hemifields. The latter was achieved by removing motion vectors above the CoM. We also included units tuned to the optic flow patterns created during translation above a ground plane:

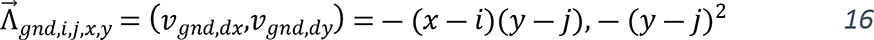

To generate valid ground templates using Eq. 16, we removed motion vectors above the CoM.

We implemented MSTd tuning to the preferred pattern using direction templates that select MT^+^ signals when they appear in appropriate spatial locations. For example, in the case of a cell tuned to radial expansion with a centrally positioned FoE, the rightward direction template pools the responses of MT+ cells tuned to rightward motion when their receptive fields coincide with the right side of the visual field. The following equations define the RF template *T*_*ψ*,*d*,*i*,*j*,*x*,*y*_ for pattern *ψ* that integrates MT^+^ cells tuned to direction *d*, normalized by the total number of pooled cells (*x̂ŷ*):

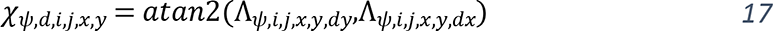

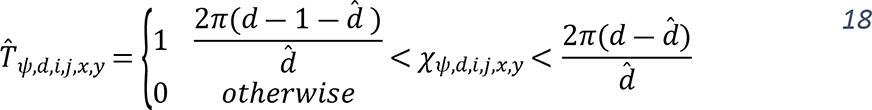

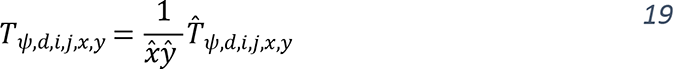

In Eq. 19, 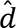 indexes the *N*^*d*^ MT preferred directions.

Within each pattern template, we sampled a small subset of the possible connections from MT^+^ units (e.g. 200 instead of 64×64×24 = 98304). Sparse connectivity is more biologically plausible and substantially improves the algorithmic efficiency of the model. We implemented this by randomly selecting only *N*^*T*^ connections to MT^+^ units within each template tuned to pattern *ψ* with CoM positioned at (*i*,*j*). Samples were drawn from a multivariate Gaussian centered on the CoM to increase the likelihood of selecting MT^+^ units positioned nearby the CoM. After drawing *N*^*T*^ pairs of (*x*,*y*) indices per template, we selected directions at each spatial location according to the uniform distribution *U*[1, *N*^*d*^]. We denote the set of *N*^*T*^ three-tuple (*x*,*y*,*d*) indices of MT^+^ units sampled within each template Ω.

The following equation matches the direction templates with the output signals from MT^+^ (*O*_*s*,*x*,*y*,*d*_) at the set of sampled spatial and directional indices (*x*,*y*,*d*) ∈ Ω to resolve the net input (*R*_*ψ*,*i*,*j*_) to the MSTd unit that prefers the pattern *ψ* with a CoM positioned at (*i*,*j*):

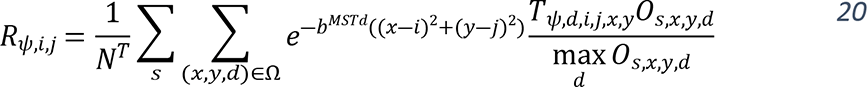

In Eq. 20, the exponential function makes MSTd units more sensitive to motion nearby the preferred CoM position and the parameter *b*^*MSTd*^ modulates how sensitivity decreases with distance. Dividing by the maximum directional input at each position ensures that dominant inputs make comparable contributions across visuotopic space before the exponential distance weighting from the CoM is applied.

### MSTd Layer 1a

We arranged MSTd units into separate layers for organizational clarity. Note that these layers do not correspond to anatomical laminae. Layer 1 corresponds to MSTd units that are driven by their pattern tuning and perform spatiotemporal averaging of their input *R*_*ψ*,*i*,*j*_. We divided the temporal (Layer 1a) and spatial (Layer 1b) averaging into two sequential steps. Layer 2 includes recurrent competitive connections that normalize the total activation while enhancing the contrast between strongly and weakly active MSTd units. We rely on the signals produced by Layer 2 for decoding the parameters specifying the observer’s self-motion.

Dynamics in Layer 1a average each MSTd unit’s net input (Eq. 21) over time:

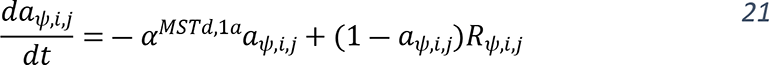

### MSTd Layer 1b

Units in MSTd Layer 1b pool the activation among units tuned to similar CoM locations and patterns along the spiral continuum (Figure S1a). We achieved this by convolving the spatial grid of signals emanating from Layer 1a units with a two-dimensional Gaussian 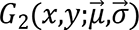

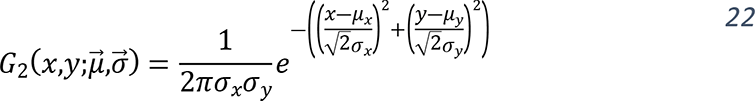

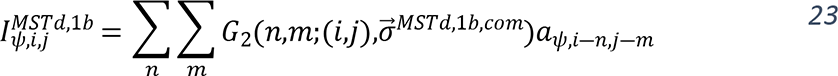

and convolving the pattern signals at each CoM position with a one-dimensional Gaussian:

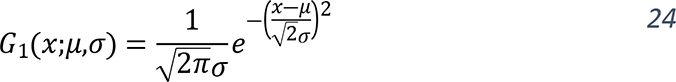

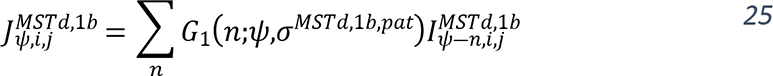

Because we arranged units tuned to similar patterns along a circular spiral pattern continuum (Figure S1b), we wrapped the convolution in Eq. 25 around the boundaries.

The dynamics of Layer 1b units obey:

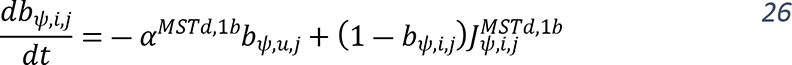

### MSTd Layer 2

The input to Layer 2 undergoes an adaptive threshold to suppress signals weaker than the mean activation value *c*_*ψ*,*i*,*j*_ achieved among units tuned to the same pattern over the recent time history

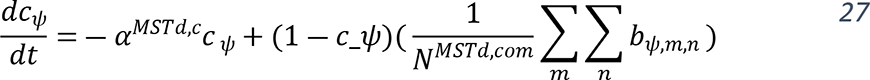

where *N*^*MSTd*,*com*^ represents the total number of units tuned to the same pattern with differing CoM positions. The net input to Layer 2 is the Layer 1b output signal with the threshold applied:

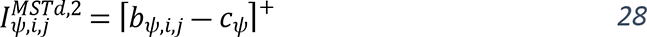

where ⌈ ⋅ ⌉^+^represents the half-wave rectification max (⋅,0).

Layer 2 integrates signals from Layer 1b and contains recurrent connections within the layer that normalize activation, facilitate competition, and stabilize signals under changing conditions (Layton & Fajen, 2016b). Units obey the recurrent competitive field with a contrast-enhancing feedback transfer function *f*(*w*) = *w*^2^, where *w* represent the unit activation (Grossberg, 1973):

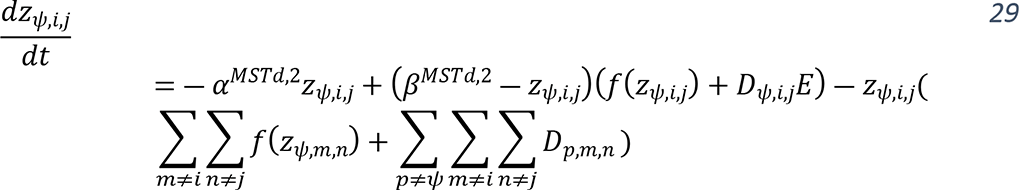

In Eq. 29, *β*^*MSTd*,2^ represents the excitatory upper bound of each unit and the terms involving *f*(⋅) implement competition among units tuned to the same pattern. The terms *D*_*ψ*,*i*,*j*_*E* and ∑_*p*≠*ψ*_ ∑_*m*≠*i*_ ∑_*n*≠*j*_ *D*_*p*,*m*,*n*_ describe excitatory and inhibitory recurrent connections, respectively, that subserve activity normalization across the layer (Boardman & Bullock, 1991). In the former term, *D*_*ψ*,*i*,*j*_ denotes the difference between each unit’s current activation and the feedforward input signal:

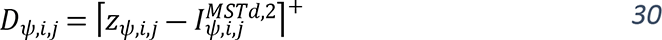

The variable *E* defines the discrepancy between input and activation across the entire layer:

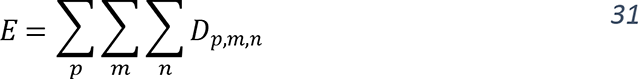

This means that the network attempts to conserve the total activation across the entire layer, not just among units tuned to the same pattern.

### Deep network decoding

We used deep learning to decode path sign, gaze offset, and path curvature during curvilinear self-motion and heading during linear self-motion. We trained separate networks to estimate each of these parameters from Layer 2 of model MSTd (Figure 12). Table 5 specifies the decoder architectures and Table 6 contains the hyperparameters used to train the networks.

**Figure 12:**
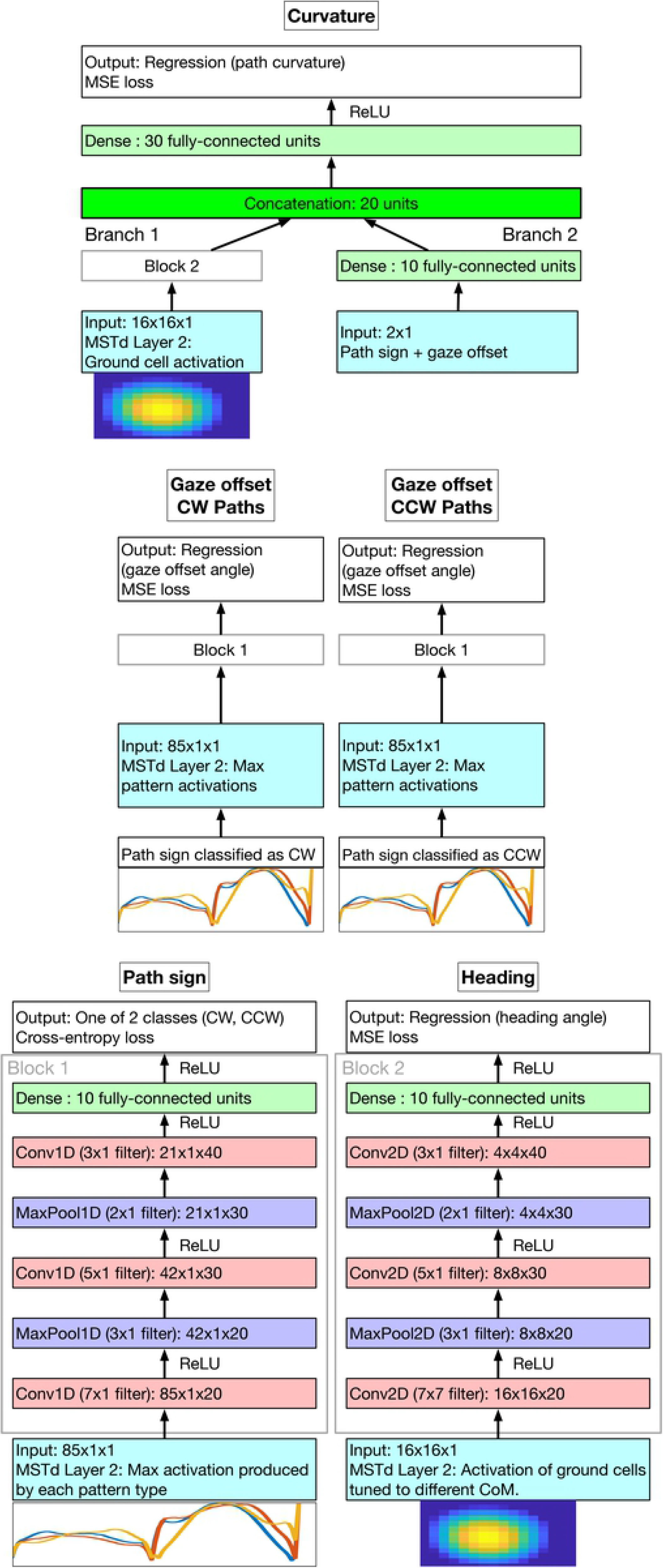
Overview of deep network decoder architectures. MSTd Layer 2 activations from the CD neural model serve as the input to the decoders. The network that classifies the CW/CCW sign of the curvilinear path (lower-left) contains interleaved 1D convolution and max pooling layers. Heading (lower-right) is decoded using a similar network, except that convolutions and max pooling operations are 2D to accommodate the spatial grid of ground unit activations. Gaze offset (center) is estimated using separate networks, conditional on the path sign prediction. Path curvature (top) is estimated using a multi-input network with convolutional (Branch 1) and fully connected (Branch 2) branches. During prediction the inputs to Branch 2 are produced by the networks that decode path sign and gaze offset.

**Table 5:** Architectures of deep network decoders. All convolutions use stride of 1 and ‘same’ padding. Max pooling operations use strides of 2.

| NETWORK | Description | Layers |
| --- | --- | --- |
| Path sign | Convolutional neural network (classification)<br>Input samples: (85x1) vectors of the max activation garnered by each MSTd Layer 2 pattern population across CoM positions.<br>Output: Binary classification of path sign. | 0: Input layer (85x1x1)<br>1: 20 7x1 filters, ReLU<br>2: 3x1 max pooling<br>3: 30 5x1 filters, ReLU<br>4: 2x1 max pooling<br>5: 40 3x1 filters, ReLU<br>6: 10 dense units, ReLU<br>7: 2 classification units, softmax, cross-entropy loss |
| Heading | Convolutional neural network (regression)<br>Input samples: (16x16) arrays of activation garnered by MSTd Layer 2 cells tuned to ground flow patterns.<br>Output: Heading angle (°) | 0: Input layer (16x16x1)<br>1-6: Same as path sign network<br>7: Regression layer, MSE loss |
| Gaze offset | 2x Convolutional neural networks (regression)<br>One network trained on gaze offsets for CW paths, the other trained on gaze offsets for CCW paths.<br>Input samples: (84x1) vectors of the max activation garnered by each MSTd Layer 2 pattern population across CoM positions.<br>Output: Gaze angle offset from straight-ahead (°) | 0-6: Same as path sign network<br>7: Regression layer, MSE loss |
| Curvature | Multi-input neural network (regression) | Branch 1: |
|  | Input branch 1 (dense): 2x1 (path sign, gaze offset angle).<br>Input branch 2 (conv): 16x16 (MSTd Layer 2 activation ground flow cells tuned to different CoM positions).<br>Output: Path curvature (1/m) | 0: Input layer (16x16x1)<br>1-6: Same as path sign network<br>Branch 2:<br>0: Input layer (2x1)<br>1: 10 dense units<br>Merge branches:<br>Concatenation layer (20x1 dense)<br>30 dense units, ReLU<br>Regression layer, MSE loss |

**Table 6:** Hyperparameters used to train deep learning network decoders.

| Hyperparameter | Value |
| --- | --- |
| Optimizer | Adam |
| Training epochs | 10,000 with early stopping (patience: 5) |
| Shuffling | Every epoch |
| Validation set | 20% of training samples. Validation checked every 50 epochs. |
| Learning rate | 1e-3 |
| Mini-batch size | 128 |
| L2 regularization | 1e-4 |

The network that classifies path sign is a 7-layer convolutional neural network (CNN) that takes as input a vector of the activations garnered by the maximally active cells tuned to each pattern type (Figure S1b). The max operation removes information about the CoM position and the vector of max input activations can be viewed as the evidence for the presence of each template pattern in the optic flow sample. Responses of many MSTd neurons have been shown to exhibit similar invariance about the position of the CoM (Graziano et al., 1994). We normalized each MSTd activity sample to values between 0 and 1 by dividing by the maximum across the pattern distribution and centered the data prior to training. The network that estimates heading is similar to the path sign network, except that it performs regression rather than classification and input features are the activations of cells tuned to ground flow that possess sensitivity to different FoE positions. We decoded from units tuned to ground flow due to its prevalence in the datasets. We trained two separate CNNs to estimate gaze offset: one specialized for CW paths, the other for CCW paths.

We estimate path curvature using a multi-input network that learns based on the ground cell features (Branch 1) and contextual information afforded by the gaze offset angle and path sign (Branch 2), both of which qualitatively alter the optic flow field. We selected ground cell features because self-motion along a curvilinear path spatially biases their activation in proportion to path curvature (Figure S1c). Gaze offset modulates the extent of the bias: gaze directed outside the future path linearizes the optic flow and gaze directed inside the future path increases the apparent curvature of the optic flow (Layton & Browning, 2014) (Li & Cheng, 2011). We transformed the labels from path radius to curvature space (Eq. 1) prior to training.

The predictions made by the decoders have hierarchical structure due to dependencies between the input features used in several networks and the parameters estimated by others. The gaze offset angle prediction is conditional on whether the path sign network predicts a CW (uses CW gaze offset network) or CCW path (uses CCW gaze offset network). Path curvature predictions depend on those made by both the path sign and gaze offset networks.

## Supporting information captions

**Figure S1:**
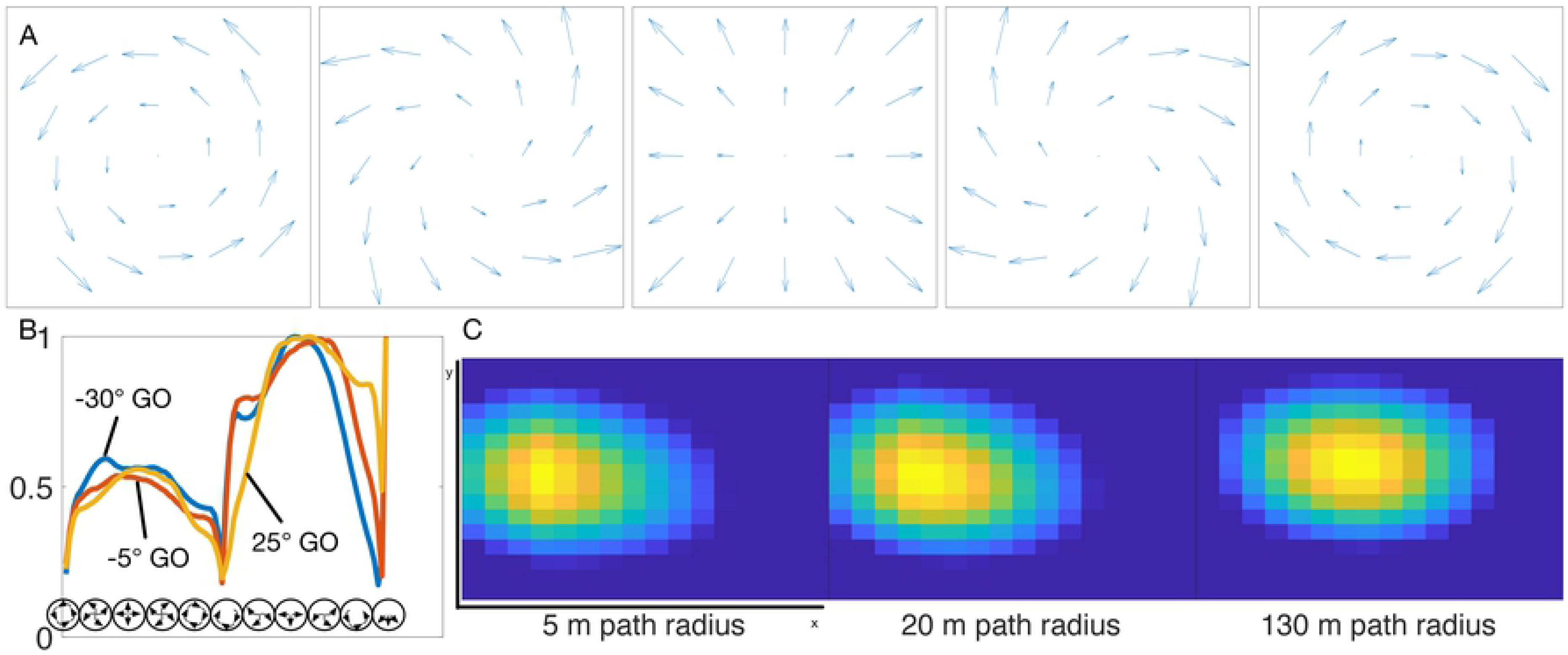
Pattern selectivity in MSTd and the effect that self-motion parameters have on population activation. (A) Sample MSTd receptive field templates spanning a continuum of patterns, from CCW centers (left), CCW spirals, radial flow, CW spirals, and CW centers (right). In each case, we simulated units tuned to CoM locations regularly spaced across the visual field in a 16×16 grid (only patterns with a central CoM are shown). (B) The max activation of MSTd units tuned to each of these patterns (left half) and hemi-field versions (right half). Units tuned to ground flow occupy the right-most position. The max is taken over units tuned to different CoM positions. The three curves show how gaze offset (GO) modulates the pattern of activation. (C) Activation of ground cells tuned to different CoM positions generated from optic flow from a 5 m (left), 20 m (center), and 130 m (right) path.

## References

Alom, M. Z., Taha, T. M., Yakopcic, C., Westberg, S., Sidike, P., Nasrin, M. S. et al. (2018). The history began from alexnet: A comprehensive survey on deep learning approaches. arXiv preprint arXiv:1803.01164.

Banks, M. S., Ehrlich, S. M., Backus, B. T., & Crowell, J. A. (1996). Estimating heading during real and simulated eye movements. Vision research, 36(3), 431–443.

Berezovskii, V. K., & Born, R. T. (2000). Specificity of projections from wide-field and local motion-processing regions within the middle temporal visual area of the owl monkey. Journal of Neuroscience, 20(3), 1157–1169.

Beyeler, M., Dutt, N., & Krichmar, J. L. (2016). 3D Visual Response Properties of MSTd Emerge from an Efficient, Sparse Population Code. J Neurosci, 36(32), 8399–8415.

Boardman, I., & Bullock, D. (1991). A neural network model of serial order recall from short-term memory.

Born, R. T., & Tootell, R. B. H. (1992). Segregation of global and local motion processing in primate middle temporal visual area. Nature, 357(6378), 497.

Britten, K. H., & van Wezel, R. J. A. (1998). Electrical microstimulation of cortical area MST biases heading perception in monkeys. Nature neuroscience, 1, 59.

Britten, K. H., & van Wezel, R. J. A. (2002). Area MST and heading perception in macaque monkeys. Cerebral cortex, 12(7), 692–701.

Browning, N. A., Grossberg, S., & Mingolla, E. (2009). A neural model of how the brain computes heading from optic flow in realistic scenes. Cogn Psychol, 59(4), 320–356.

Cameron, S., Grossberg, S., & Guenther, F. H. (1998). A self-organizing neural network architecture for navigation using optic flow. Neural Computation, 10(2), 313–352.

Cheng, J. C., & Li, L. (2012). Effects of reference objects and extra-retinal information about pursuit eye movements on curvilinear path perception from retinal flow. J Vis, 12(3).

Danz, A. D., Angelaki, D. E., & DeAngelis, G. C. (2020). The Effects of Depth Cues and Vestibular Translation Signals on the Rotation Tolerance of Heading Tuning in Macaque Area MSTd. eNeuro, 7(6).

Duffy, C. J., & Wurtz, R. H. (1991). Sensitivity of MST neurons to optic flow stimuli. I. A continuum of response selectivity to large-field stimuli. Journal of neurophysiology, 65(6), 1329–1345.

Duffy, C. J., & Wurtz, R. H. (1995). Response of monkey MST neurons to optic flow stimuli with shifted centers of motion. Journal of Neuroscience, 15(7), 5192–5208.

Ganguli, D., & Simoncelli, E. P. (2014). Efficient sensory encoding and Bayesian inference with heterogeneous neural populations. Neural Comput, 26(10), 2103–2134.

Georgopoulos, A. P., Schwartz, A. B., & Kettner, R. E. (1986). Neuronal population coding of movement direction. Science, 233(4771), 1416–1419.

Gibson, J. J. (1950). The perception of the visual world. Houghton Mifflin.

Glaser, J. I., Benjamin, A. S., Chowdhury, R. H., Perich, M. G., Miller, L. E., & Kording, K. P. (2020). Machine Learning for Neural Decoding. eNeuro, 7(4).

Graziano, M. S., Andersen, R. A., & Snowden, R. J. (1994). Tuning of MST neurons to spiral motions. J Neurosci, 14(1), 54–67.

Grossberg, S. (1973). Contour Enhancement, Short Term Memory, and Constancies in Reverberating Neural Networks. Studies in Applied Mathematics, 52, 213–257.

Grossberg, S. (1980). Intracellular mechanisms of adaptation and self-regulation in self-organizing networks: the role of chemical transducers. Bulletin of Mathematical Biology, 42(3), 365–396.

Grossberg, S. (2020). A Path Toward Explainable AI and Autonomous Adaptive Intelligence: Deep Learning, Adaptive Resonance, and Models of Perception, Emotion, and Action. Frontiers in Neurorobotics, 14.

Gu, Y., Fetsch, C. R., Adeyemo, B., Deangelis, G. C., & Angelaki, D. E. (2010). Decoding of MSTd population activity accounts for variations in the precision of heading perception. Neuron, 66(4), 596–609.

Gu, Y., Watkins, P. V., Angelaki, D. E., & DeAngelis, G. C. (2006). Visual and nonvisual contributions to three-dimensional heading selectivity in the medial superior temporal area. J Neurosci, 26(1), 73–85.

Gu, Y., DeAngelis, G. C., & Angelaki, D. E. (2012). Causal links between dorsal medial superior temporal area neurons and multisensory heading perception. Journal of Neuroscience, 32(7), 2299–2313.

Hubel, D., & Wiesel, T. (2012). David Hubel and Torsten Wiesel. Neuron, 75(2), 182–184.

Hubel, D. H. (1982). Exploration of the primary visual cortex, 1955–78. Nature, 299(5883), 515–524.

Inaba, N., Shinomoto, S., Yamane, S., Takemura, A., & Kawano, K. (2007). MST neurons code for visual motion in space independent of pursuit eye movements. J Neurophysiol, 97(5), 3473–3483.

Jazayeri, M., & Movshon, J. A. (2006). Optimal representation of sensory information by neural populations. Nat Neurosci, 9(5), 690–696.

Komatsu, H. I. D. E. H. I. K. O., & Wurtz, R. O. B. E. R. T. H. (1988). Relation of cortical areas MT and MST to pursuit eye movements. I. Localization and visual properties of neurons. Journal of neurophysiology, 60(2), 580–603.

Krizhevsky, A., Sutskever, I., & Hinton, G. E. (2012). Imagenet classification with deep convolutional neural networks. Advances in neural information processing systems, 25, 1097–1105.

Layton, O. W., & Fajen, B. R. (2016a). A Neural Model of MST and MT Explains Perceived Object Motion during Self-Motion. J Neurosci, 36(31), 8093–8102.

Layton, O. W., & Fajen, B. R. (2017). Possible role for recurrent interactions between expansion and contraction cells in MSTd during self-motion perception in dynamic environments. J Vis, 17(5), 5.

Layton, O. W., & Browning, N. A. (2012). Recurrent competition explains temporal effects of attention in MSTd. Frontiers in computational neuroscience, 6, 80.

Layton, O. W., & Fajen, B. R. (2016b). Competitive dynamics in MSTd: A mechanism for robust heading perception based on optic flow. PLoS computational biology, 12(6), e1004942.

Layton, O. W., Mingolla, E., & Browning, N. A. (2012). A motion pooling model of visually guided navigation explains human behavior in the presence of independently moving objects. Journal of vision, 12(1), 20–20.

Layton, O. W., & Niehorster, D. C. (2019). A model of how depth facilitates scene-relative object motion perception. PLoS computational biology, 15(11), e1007397.

Layton, O. W., & Browning, N. A. (2014). A unified model of heading and path perception in primate MSTd. PLoS Comput Biol, 10(2), e1003476.

Layton, O. W., & Fajen, B. R. (2020). Computational Mechanisms for Perceptual Stability using Disparity and Motion Parallax. J Neurosci, 40(5), 996–1014.

Li, L., & Cheng, J. C. K. (2011). Perceiving path from optic flow. Journal of Vision, 11(1), 22–22.

Logan, D. J., & Duffy, C. J. (2006). Cortical area MSTd combines visual cues to represent 3-D self-movement. Cereb Cortex, 16(10), 1494–1507.

Longuet-Higgins, H. C., & Prazdny, K. (1980). The interpretation of a moving retinal image. Proc. R. Soc. Lond. B, 208(1173), 385–397.

Maunsell, J. H., & van Essen, D. C. (1983). The connections of the middle temporal visual area (MT) and their relationship to a cortical hierarchy in the macaque monkey. Journal of Neuroscience, 3(12), 2563–2586.

Mikami, A., Newsome, W. T., & Wurtz, R. H. (1986). Motion selectivity in macaque visual cortex. II. Spatiotemporal range of directional interactions in MT and V1. J Neurophysiol, 55(6), 1328–1339.

Mineault, P. J., Khawaja, F. A., Butts, D. A., & Pack, C. C. (2012). Hierarchical processing of complex motion along the primate dorsal visual pathway. Proc Natl Acad Sci U S A, 109(16), E972–80.

Orban, G. A., Lagae, L., Verri, A., Raiguel, S., Xiao, D., Maes, H. et al. (1992). First-order analysis of optical flow in monkey brain. Proceedings of the National Academy of Sciences, 89(7), 2595–2599.

Page, W. K., & Duffy, C. J. (1999). MST neuronal responses to heading direction during pursuit eye movements. Journal of neurophysiology, 81(2), 596–610.

Perrone, J. A. (2018). Visual-vestibular estimation of the body’s curvilinear motion through the world: A computational model. J Vis, 18(4), 1.

Perrone, J. A., & Krauzlis, R. J. (2008). Vector subtraction using visual and extraretinal motion signals: a new look at efference copy and corollary discharge theories. J Vis, 8(14), 24.1–2414.

Perrone, J. A. (1992). Model for the computation of self-motion in biological systems. JOSA A, 9(2), 177–194.

Raudies, F., & Neumann, H. (2013). Modeling heading and path perception from optic flow in the case of independently moving objects. Front Behav Neurosci, 7, 23.

Raudies, F., & Neumann, H. (2012). A review and evaluation of methods estimating ego-motion. Computer Vision and Image Understanding, 116(5), 606–633.

Royden, C. S. (2002). Computing heading in the presence of moving objects: a model that uses motion-opponent operators. Vision research, 42(28), 3043–3058.

Royden, C. S., Crowell, J. A., & Banks, M. S. (1994). Estimating heading during eye movements. Vision research, 34(23), 3197–3214.

Saito, H.-a., Yukie, M., Tanaka, K., Hikosaka, K., Fukada, Y., & Iwai, E. (1986). Integration of direction signals of image motion in the superior temporal sulcus of the macaque monkey. Journal of Neuroscience, 6(1), 145–157.

Sanger, T. D. (1996). Probability density estimation for the interpretation of neural population codes. Journal of neurophysiology, 76(4), 2790–2793.

Serre, T. (2019). Deep learning: the good, the bad, and the ugly. Annual Review of Vision Science, 5, 399–426.

Shah, S., Dey, D., Lovett, C., & Kapoor, A. (2017). AirSim: High-Fidelity Visual and Physical Simulation for Autonomous Vehicles. arXiv, 1705.05065v2.

Shenoy, K. V., Bradley, D. C., & Andersen, R. A. (1999). Influence of gaze rotation on the visual response of primate MSTd neurons. Journal of Neurophysiology, 81(6), 2764–2786.

Simonyan, K., & Zisserman, A. (2014). Very Deep Convolutional Networks for Large-Scale Image Recognition. arXiv, 1409.1556v6.

Steinmetz, S. T., Layton, O. W., Powell, N. V., & Fajen, B. R. (2021). A Dynamic Efficient Sensory Encoding Approach to Adaptive Tuning in Neural Models of Visual Motion Processing. https://www.biorxiv.org/content/10.1101/2021.06.17.448823v1.abstract

Tanaka, K., Hikosaka, K., Saito, H.-a., Yukie, M., Fukada, Y., & Iwai, E. (1986). Analysis of local and wide-field movements in the superior temporal visual areas of the macaque monkey. Journal of Neuroscience, 6(1), 134–144.

Tanaka, K. E. I. J. I., Sugita, Y. O. I. C. H. I., Moriya, M. A. D. O. K. A., & Saito, H.-a. (1993). Analysis of object motion in the ventral part of the medial superior temporal area of the macaque visual cortex. Journal of Neurophysiology, 69(1), 128–142.

Warren, W. H., & Hannon, D. J. (1990). Eye movements and optical flow. JOSA A, 7(1), 160–169.

Warren, W. H., Mestre, D. R., Blackwell, A. W., & Morris, M. W. (1991). Perception of circular heading from optical flow. Journal of Experimental Psychology: Human Perception and Performance, 17(1), 28.

Warren, W. H., Morris, M. W., & Kalish, M. (1988). Perception of translational heading from optical flow. Journal of Experimental Psychology: Human Perception and Performance, 14(4), 646.

Warren, W. H., & Saunders, J. A. (1995). Perceiving heading in the presence of moving objects. Perception, 24(3), 315–331.

Weinzaepfel, P., Revaud, J., Harchaoui, Z., & Schmid, C. (2013). DeepFlow: Large Displacement Optical Flow with Deep Matching. Proceedings from 2013 IEEE International Conference on Computer Vision.

Xu, H., Wallisch, P., & Bradley, D. C. (2014). Spiral motion selective neurons in area MSTd contribute to judgments of heading. J Neurophysiol, 111(11), 2332–2342.

Yu, X., Hou, H., Spillmann, L., & Gu, Y. (2018). Causal evidence of motion signals in macaque middle temporal area weighted-pooled for global heading perception. Cerebral Cortex, 28(2), 612–624.

Zhou, B., Krähenbühl, P., & Koltun, V. (2019). Does computer vision matter for action. Sci Robot, 4(30).

